# Continuous target-specific mutagenesis and rapid gene evolution by diversity-generating retroelements in *Escherichia coli*

**DOI:** 10.64898/2026.07.21.739853

**Authors:** Yang Liu, Yangqi Gu, Ganesh Agam, Fabian Rehm, Martin Spinck, Jason Chin, Philipp Holliger

**Affiliations:** MRC Laboratory of Molecular Biology, Francis Crick Avenue, Cambridge CB2 0QH, UK

## Abstract

Sequence-programmable directed evolution systems have great potential to accelerate bioengineering. Diversity-generating retroelements (DGRs) are natural hypermutation systems widely distributed in prokaryotes and bacteriophages with the capacity to introduce diverse mutations at template-specified sites of target genes. Here, we show that DGRs can be installed in *E. coli* and reprogrammed for the continuous, iterative mutagenesis of user-defined target genes. We show that the DGR template RNA can be reprogrammed for gene- and residue-specific mutagenesis, leaving untargeted, adjacent residues unchanged. Furthermore, we establish continuous DGR-enabled mutagenesis with conjugation-mediated horizontal gene transfer of target genes (HGT-DGR) into a new host for the progressive accumulation of target-specific mutations. Iterative HGT-DGR mutagenesis over seven cycles yielded an average mutation load of approximately 6% across adenine positions in the target segment, generating a diverse library of variants comprising 40% mutant sequences, with a median pairwise Hamming distance of 4 among mutant variants. HGT-DGR enables iterative diversification of either the same or different user-specified segments of the target gene, as demonstrated with the directed evolution of the *M. mazei* pyrrolysyl-tRNA synthetase for non-canonical amino acid incorporation. HGT-DGR provides a simple, low-cost, sequence-programmable system that enables iterative, position-specific and tunable in vivo mutagenesis of any target sequence for applications in biotechnology and medicine.

## Introduction

Evolutionary adaptation at the molecular level is based on the process of diversification (mutation) coupled with selection to yield biomolecules with new or improved functions. Directed evolution seeks to leverage these principles in an accelerated and targeted manner (*1*). However, despite great successes in harnessing evolution for bioengineering, further improvements in speed, efficiency and programmability are desirable, especially for applications in biomedicine and biotechnology. Traditionally, directed evolution approaches have utilised *in vitro* methods for diversity generation such as error-prone PCR, which randomly introduces mutations across a defined amplicon, and site-saturation mutagenesis, where sites of interest are specifically randomized. Following a round of diversification, gene variants are transformed into cells (or viruses) where they are subjected to screening or selection. Enriched gene variants serve as templates for subsequent rounds of *in vitro* diversification, transformation and selection. While these approaches allow a great deal of control over mutation and selection parameters, iterative cycles of mutagenesis can be slow, costly and inefficient as they only explore a limited area of the sequence and phenotype space of any target. This has motivated the development of *in vivo* evolution approaches for the continuous mutation and selection. *In vivo* hypermutation is regarded as a key feature of continuous evolution systems, because it eliminates the need to repeatedly generate mutants *in vitro* and thereby enables a more rapid, autonomous, natural-evolution-like process in the laboratory (*2, 3*). In practice, the workflow can be run in a continuous or semi-continuous mode by implementing screening/selection with different levels of automation.

Early examples include the use of mutator strains in combination with phage display (*4*). This was followed by the development of phage-assisted continuous evolution (PACE) (*5*). PACE together with a mutator plasmid (*6*) enables hundreds of rounds of *in vivo* mutagenesis and selection without human intervention, which can greatly accelerate the speed of discovery. However, while these approaches are rapid, and allow exploration of a much wider sequence space, there is limited control over mutation parameters and target sequences. This has led to the development of a variety of strategies that allow a more targeted approach to *in vivo* mutagenesis. These include orthogonal episomes in both yeast (*K. lactis*) and bacteria (*E. coli*) that are replicated by a dedicated, error-prone DNA polymerase that does not participate in genome replication (*7–10*); targeted localization of promiscuous base deaminases via their fusion to a helicase or a T7 RNA polymerase (*11–15*); nCas9-mediated recruitment of an error-prone DNA polymerase (*16*); or via the use of a reverse transcriptase to generate ssDNA for targeted recombination where the mutagenesis occurs by reverse-transcription or via a deaminase fused to the reverse transcriptase (*17–19*). However, all of these approaches are limited to various extents in terms of their mutational target windows, mutational spectra, and degree of orthogonality with respect to genome replication. Importantly, these continuous evolution strategies do not allow mutagenesis to be programmed at single-nucleotide resolution across user-defined positions. There is therefore a need for sustainable *in vivo* approaches for site-saturation mutagenesis of user-defined positions, an advance that would enable more efficient, (structure-)guided targeting of relevant regions in evolved molecules.

Retron editors have been leveraged for targeted mutagenesis (*20–23*) and have recently been adapted for continuous site-specific saturation mutagenesis (*24*) via efficient conjugation mediated transfer of target genes between host carrying retron libraries. This enabled *in vivo* mutagenesis and continuous iteration of the conjugation-editing cycles with selection. In this approach, the library diversity emerged multiplicatively and was shown to enable the evolution of aminoacyl-tRNA synthetase (aaRS) variants that selectively incorporate non-canonical amino acids (ncAAs).

Here we describe an alternative approach for sustainable, iterative, residue-specific mutagenesis of user-defined target sequences *in vivo*, leveraging the properties of diversity-generating retroelements (DGRs). DGRs are widespread in prokaryotes and bacteriophages and serve to hypermutate specific gene regions. In the prototypical DGR from the *Bordetella* bacteriophage, diversification of the bacteriophage’s receptor-binding protein enables adaptation to the loss of potential surface receptors by its host *Bordetella* (*25*). Strikingly, natural DGR-mediated diversity significantly exceeds the 10^14-16^ variation of the vertebrate immune system (*26*), achieving as much as 10^20-30^ variation for a given DGR-variable protein (*27, 28*). DGRs consist of three essential components: 1) a target gene segment known as the variable region (VR); 2) a DNA template region (TR), which can be transcribed into TR-RNA and used as a template for reverse transcription; 3) a unique reverse transcriptase that can specifically introduce random bases exclusively at adenine positions. Reverse-transcribed and adenine-mutagenized cDNA can then be integrated into the VR via a retrohoming mechanism.

To harness the residue-specific mutagenesis enabled by DGRs in *E. coli* which lacks DGRs, we engineered a synthetic retrohoming mechanism in *E. coli* and demonstrated that it can be reprogrammed to precisely mutagenize defined target sequences with near-perfect residue specificity, as encoded by adenines in a user-defined template region (TR). Indeed, a similar DGR-based diversification strategy was also reported recently (*19*).

However, unlike natural DGR systems, existing implementations in *E. coli* are not readily sustainable over extended periods because host cells poorly tolerate prolonged recombination stress, while mutagenesis of the TR itself destabilizes the sequence space explored during evolution. This prevents the use of DGR mutagenesis for continuous evolution. To overcome this, we addressed both DGR instability and TR mutagenesis by coupling DGR diversification to iterative horizontal gene transfer (HGT) via bacterial conjugation. This enabled stable and sustainable long-term operation of DGRs in *E. coli* enabling the cumulative accumulation of edited variants up to 40% in the cultured population within one week. Finally, we demonstrate *in vivo* mutagenesis at two distinct loci to rapidly evolve aaRS variants with altered substrate specificity.

## Results

### Implementing the BPP1 DGR system in *E. coli*

To establish a DGR-based hypermutation system in *E. coli*, which does not naturally harbour a DGR, we focused on the prototypical DGR system from the *Bordetella*-infecting BPP1 bacteriophage. We first expressed the natural components of the BPP1-DGR system, including the reverse transcriptase (bRT), accessory variability determinant protein (Avd), the wild-type (WT) VR and the WT TR (*29*). Given that the mechanism of DGR retrohoming remains unknown and does not naturally operate in *E. coli*, we independently developed a synthetic strategy to mimic the DGR retrohoming process, in parallel with another recently reported study (*19*). In brief, we constructed a cassette expressing the highly-efficient single-stranded DNA (ssDNA) annealing protein CspRecT from *Collinsella stercoris* and a dominant negative mutant of MutL (mMutL (E32K)) **(Fig. 1A)** (*30, 31*). CspRecT has been widely used to mediate the integration of ssDNA by facilitating annealing at plasmid or genome replication forks (*30, 31*). Previous studies had implicated mismatch repair system (MMR) in the retrohoming process of DGRs given that many MutS-like homologs are found in the natural DGR systems (*28*). We therefore utilized mMutL to facilitate suppression of the *E. coli* MMR system **(Fig. 1A)** (*32*). During the course of this study, mMutL has also been independently reported to enhance recombineering efficiency via suppression of mismatch repair, including in the context of DGR based mutagenesis in *E. coli* (*19*).

**Figure 1.**
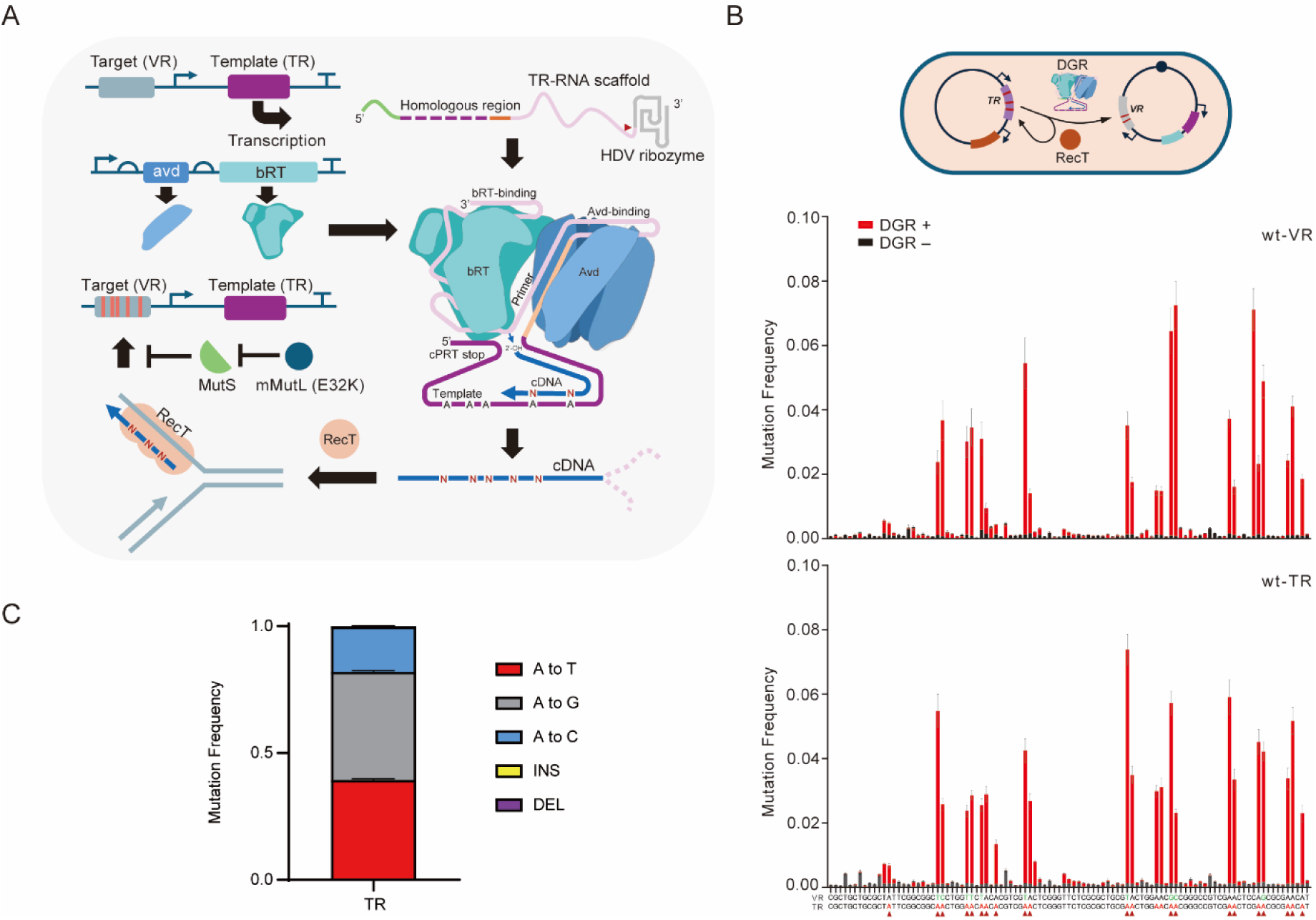
Implementing diversity-generating retroelements (DGR) in E. coli. **A,** Design of the DGR circuit, which contains 3 different components: 1) Template RNA, (TR) which is excised into a mature TR scaffold with a self-cleaving HDV ribozyme. 2) DGR reverse transcription cassette including the reverse transcriptase (bRT) and assisting protein (Avd). 3 CspRecT and mMutL (E32K) for homologous recombination. **B,** Schematic of engineered homing mechanism for DGR in E. coli and 24h mutation frequency barplots of VR and TR. The VR and the TR sequence is shown at the bottom of the barplot with Adenosine positions highlighted in red with triangle and the different sequence between TR and VR is highlighted in green. DGR+: With DGR. DGR-: Without DGR. **C,** Mutational spectrum of DGR based on the TR mutations. Error bars, s.d. (n = 3)

We quantified mutation rate in DGR-mediated mutagenesis using next-generation sequencing (NGS). In our 2-plasmid system, the mutagenesis in 24h of the WT VR via the WT TR revealed DGR-specific mutations with above background mutagenesis only observed for adenine positions as expected for DGR retrohoming mediated mutagenesis **(Fig. 1B, S1).**

It is important to note that mutation numbers do not scale linearly with DNA replication or cell division. This is because in the DGR system mutations originate from both reverse transcription and recombination, and the rates of these processes are related to the intracellular levels of the enzymes involved. It is well known that intracellular enzyme levels fluctuate throughout a growth cycle, and the steady state period is short and not parallel to the cell growth rate (*33*). We also conducted a test and found that altering the times of cell divisions throughout the DGR induction by controlling the initial cell number (diluting the overnight culture 100-fold or 1000-fold) did not significantly change the mutation frequency on VR target sequence after a single round of DGR induction **(Fig. S2)**. We therefore define the mutation efficiency as the mean mutation frequency at all adenine positions within the target segment, as measured over a defined time interval. Because the VR sequence differs from the TR, new VR variants can arise either from direct TR to VR replacement (recombination) or from DGR-mediated adenine mutagenesis. Therefore, when calculating the VR mutation frequency strictly attributable to DGR activity, we restricted the analysis to positions at which both the VR and TR encode adenine in the unedited reference sequences. Using these criteria, we obtained an average mutation efficiency of 3.54% ± 1.58% for TR in 24h, and 1.85% ± 1.08% for VR under the same condition **(Fig. S3)**.

Next, we tested alternative TR sequence designs: one retaining 300 bp of the WT TR upstream sequence and the other retaining only 21 bp. According to previous research, the TR upstream sequence encodes the 3’ end of the cDNA and is involved in the termination of reverse transcription **(Fig. S4A)** (*34*). We hypothesized that 21 bp would be sufficient to terminate the reverse transcription (*35*). Indeed, we found that the shorter TR is sufficient to accommodate DGR-like mutations with higher mutation frequencies compared to the longer TR design **(Fig. S4B, C)**. The mutational spectrum at adenine positions was consistent with previous studies (*36*), with about 2.2 – 2.4 fold higher mutational bias towards Guanine or Thymine compared to Cytosine with negligible insertion or deletions (indels) **(Fig. 1C)**.

We found that both CspRecT and mMutL are indispensable for DGR-based retrohoming mutagenesis **(Fig. S5)**. Notably, this synthetic retrohoming strategy not only rewrites VR but also significantly mutates TR, continuously erasing adenine from TR **(Fig. 1B, S6)**. In our initial design, the mutation rate of the TR was higher than that of the VR **(Fig. S7)**. These undesirable TR mutations progressively restrict the capacity of the system for sustained mutagenesis. We attempted to increase the VR mutation rate by testing different orientations of VR in respect to the replication fork, since previous studies showed that CspRecT favors the integration of ssDNA on the lagging strand over the leading strand (*37*). Indeed, we observed an increased (1.35-fold) VR mutation rate when targeting the lagging strand, as expected **(Fig. S8)**.

### Reprogramming the BPP1 DGR system to target a gene of interest

Next, we extracted 150 VR sequences of BPP1-like DGR systems from a public database (*38*). Multi-sequence alignment of the VR protein sequences from BPP1-like DGRs showed high diversity, which suggests that the VR can target a wide range of sequences **(Fig. S9A)**. On the basis of this analysis and previous attempts to reprogram the DGR system (*19, 39*), we sought to demonstrate retargeting of the DGR system for user-defined sequence mutagenesis.

To this end, we reprogramed the BPP-1 DGR system with the superfolder green fluorescent protein (*sfgfp*) gene with an *ochre* codon at T65 as the target and designed a TR targeting the codons spanning from C48 - M78. If the DGR system is successfully reprogrammed, we should observe adenine-specific diversification within the range of codons C48 - M78 (bases 124 - 216) on the *sfgfp* gene. After inducing the DGR system, the synthetic retrohoming components, and harbouring both the reprogrammed TR and VR, we indeed observed DGR-dependent mutagenesis of the targeted *sfgfp* gene region **(Fig. S9B)**. Interestingly, when designing the TR corresponding to position T65 (ACN) as a tri-adenine (AAA), NGS revealed that DGR generated a diverse set of codons at this site, 24% of reads led to nonsense suppression, and 0.016% of reads showed reversion mutation **(Fig. S9C)**. In a 12h DGR induction test, fluorescence-activated cell sorting (FACS) enriched multiple sfGFP variants, increasing the fluorescence intensity of sfGFP by up to 55% **(Fig. S10)**. Thus, we could readily reprogram the BPP-1 DGR system to mutagenize a defined region of a specified target gene, and implement effective lab evolution.

### Design principles for an efficient DGR-based evolution system in *E. coli*

The above experiments indicated that the mutation efficiencies of the reprogrammed TR and VR differed from those of the natural VR and TR. Because multiple parameters may affect the efficiency of a synthetic DGR system in *E. coli*, we next explored different designs to derive a set of general design principles.

First, the function of our DGR system in *E. coli* is dependent on the efficiency of CspRecT-mediated recombination: we hypothesized that the length of the homology arm between the single-stranded cDNA and the target may critically affect the efficiency of recombination and thereby the fixation of the RT-induced mutations. Considering that the positions of mutations in the DGR cDNA are scattered, we consider the entire TR as the homology arm. Therefore, to test the effect of homology arm length on the mutation rate of the VR, we designed TRs with different lengths in the same region of *sfgfp*. We found that longer TRs (93 bp) significantly increased the frequency of DGR-characteristic VR mutations in 24h. Thus, mutation frequency can be tuned to some extent by the length of the TR sequence **(Fig. 2A)**. Next, we hypothesized that the efficiency of DGR-mediated mutagenesis might be influenced by the intracellular cDNA concentration as the cDNA produced by TR reverse transcription might be degraded by various host DNA exonucleases. Indeed, it had previously been shown that an exonuclease knock-out strain could improve the efficiency of retron recombineering (*21, 40*). Therefore, to achieve higher intracellular cDNA concentrations, we used an *E. coli* Δ*sbcB*, Δ*recJ* and Δ*exoX* deletion strain in which three exonuclease genes are deleted from the genome (*24, 41*). Indeed, combining optimal TR length (93 bp) with the triple knockout strain nearly doubled average mutation frequencies in the DGR system. Thus, higher DGR mutation frequencies can be obtained by TR extension and inhibition of intracellular cDNA degradation **(Fig. 2B)**.

**Figure 2.**
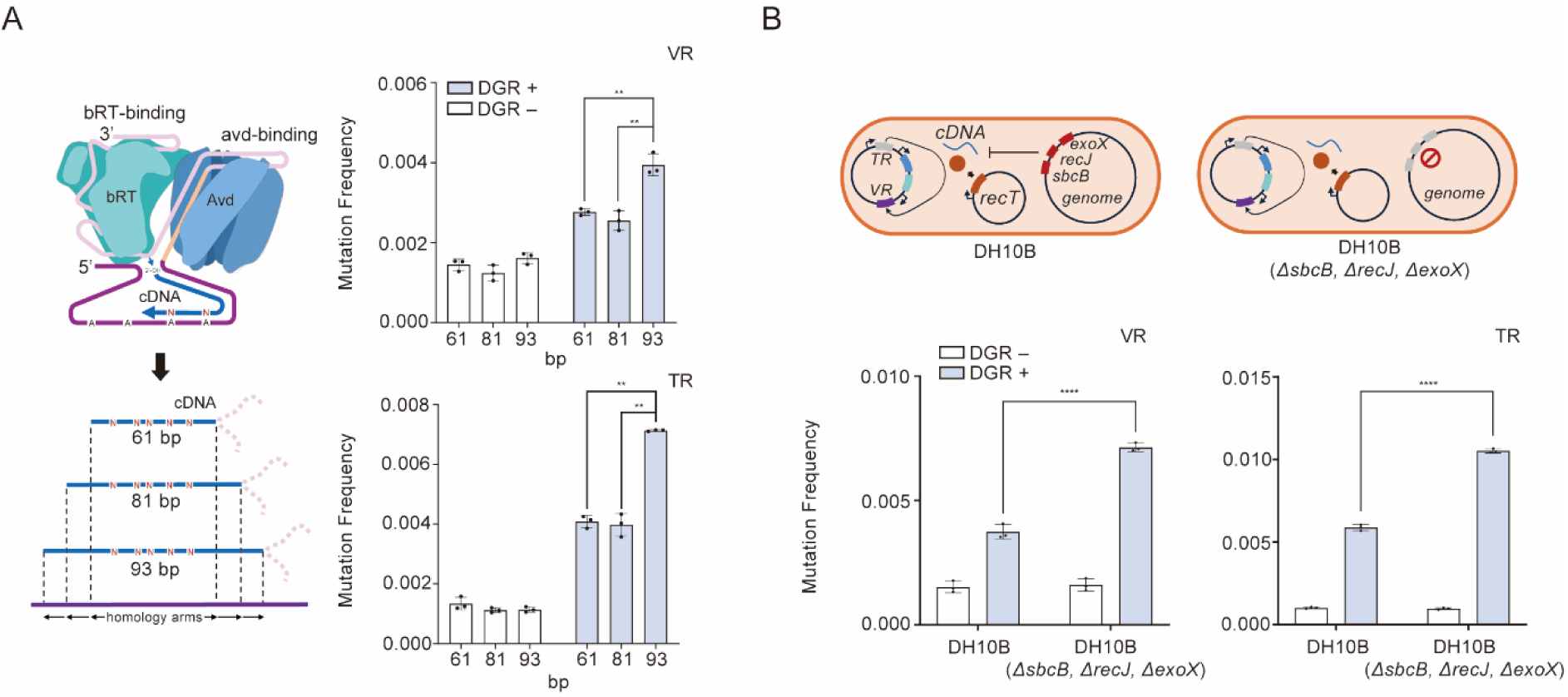
Optimization of DGR-mediated mutagenesis in E. coli. **A,** Increasing the length of homology arms for recombination increases the 24h mutational frequency for both VR and TR. Statistical difference was determined by a two-tailed unpaired t test: (VR, 61/93), p = 0.0021, t = 7.085, (VR, 81/93), p = 0.0029, t = 6.512, and a two-tailed Welch’s t test: (VR, 61/93), p = 0.0013, t = 25.70, (TR, 81/93), p = 0.0047, t = 14.30. **B,** Using a strain with genomically deleted exonucleases SbcB, RecJ and ExoX increases the 20h mutational frequency. Statistical difference was determined by a two-tailed unpaired t test for VR, p<0.0001, t = 17.20, and for TR, p<0.0001, t = 36.36. p value summary: ****p value < 0.0001, 0.0001 < ***p value < 0.001, 0.001 < **p value < 0.01, 0.01 < *p value < 0.05, p value ≥ 0.05: n.s. Error bars, s.d. (n = 3)

### Renewing TR and host via HGT enables sustainable DGR-mediated hypermutation

Having established DGR-mediated hypermutation in *E. coli* we next sought to address two constraints that limit its application for continuous directed evolution. First, the sequence space explored by DGR diversification is strictly specified by the template region (TR) and the sequence space that can be searched is limited by a finite culture volume. For applications that require exploration of a large sequence space, the current mutation rate may therefore generate too few variants per unit time to achieve meaningful coverage. Therefore, prolonged cultivation and repeated rounds of screening/selection are required. Under these conditions, TR integrity becomes essential to keep mutagenesis going and the desired search space constant across time. However, we and others have found that in a synthetic DGR implementation that the TR itself is mutated (*19*), causing drift of the encoded target sequence, which progressively reduces mutagenesis efficiency by preferentially eliminating dA from the TR sequence. Second, expression of RecT and mMutL imposes a substantial growth burden on the host cell providing a selective pressure to attenuate or revert to wild-type, leading to the progressive loss of functional DGR genotypes at the population level. Indeed, when we serially subcultured a strain carrying the DGR system, we observed that the DGR-mediated mutation frequency at population level ceased to increase and even dropped by the third day of culture **(Fig. S11)**.

We hypothesized that both limitations could be overcome by moving the VR target to fresh host cells carrying the desired TR using horizontal gene transfer (HGT). This process would simultaneously renew both the host genetic background and the TR, while allowing mutations to iteratively accumulate within the VR **(Fig. 3A)**.

**Figure 3.**
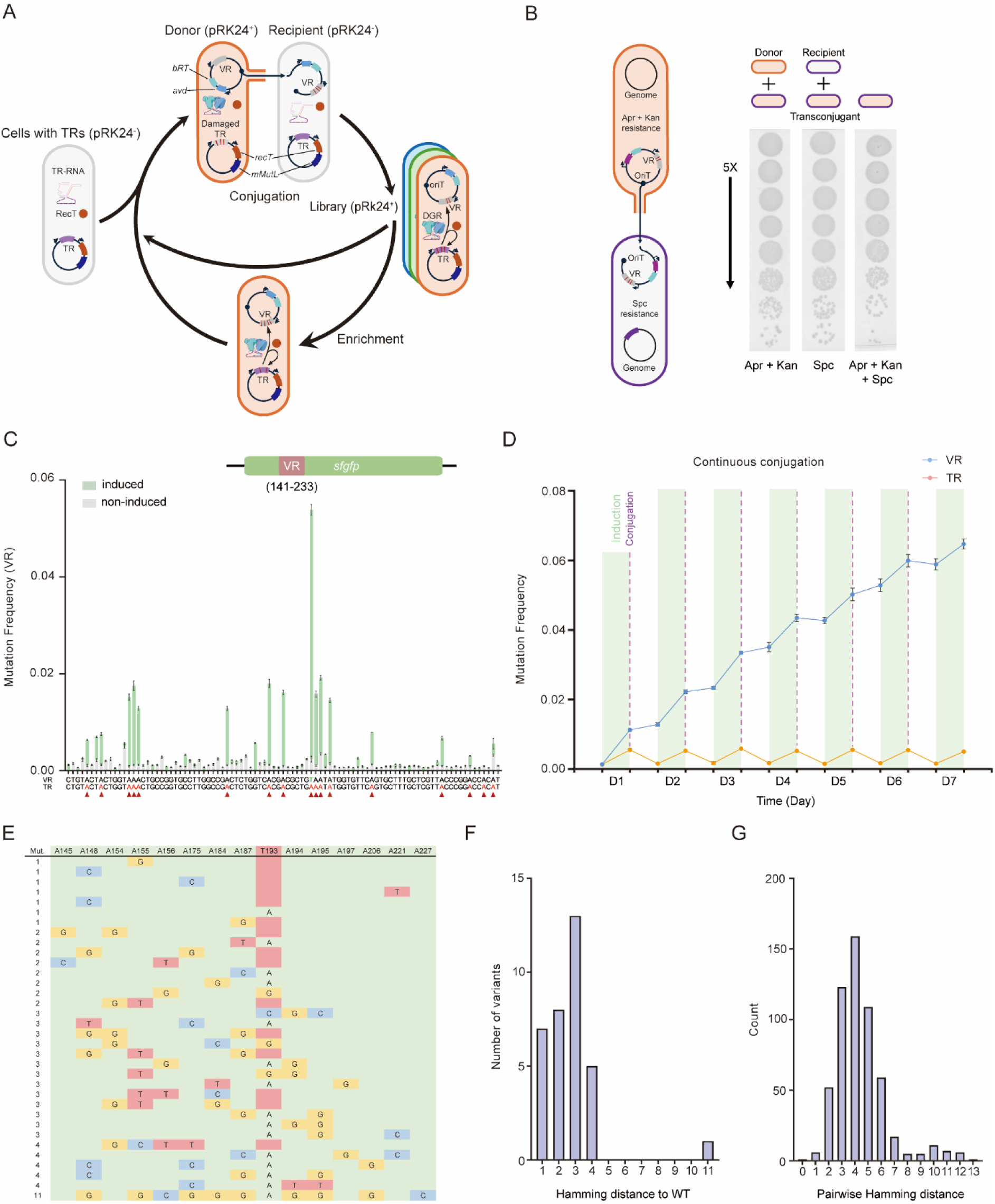
Horizontal gene transfer mediated DGR evolution via conjugation. **A,** Schematic of HGT-DGR, where the sfgfp, brt and avd genes are incorporated onto a transferrable conjugative plasmid (pRK24+) with origin of transfer (OriT) and conjugation into recipient cells containing no conjugative plasmid (pRK24-) enables the high-fidelity transfer of the VR library. **B,** Spotting assay confirmed the high transfer rate of brt, avd, sfgfp genes-containing conjugative plasmid into empty cells. The conjugative plasmid harbours resistance genes for both kanamycin and apramycin, while the recipient contains a genomically integrated spectinomycin resistance gene. **C,** Mutation frequency (10h) barplot of VR (sfGFP) on the conjugative plasmid. Adenosine positions are highlighted in red with triangle and the sequence differences between TR and VR are highlighted in green. The thymine site T193 in VR, which does not match the TR sequence is not included in the calculation. **D,** Averaged mutation frequency (10h) of adenine position from 7 days of continuous conjugation. **E,** Adenine site mutation profile of single-colony sequenced variants after 7 days of continuous hypermutation. **F,** Distribution of Hamming distances to the WT fingerprint for 34 mutant VR variants isolated after 7 days, showing the number of differing positions per variant. **G,** Distribution of pairwise Hamming distances among the same 34 fingerprints (n = 561 pairwise comparisons), reporting the number of differing positions between variant pairs (median 4; IQR 3–5). Error bars, s.d. (n = 3)

We initially tested whether the constant DGR system components (*avd* and *brt*) together with the sfgfp (VR) **(Fig. S9B)** could be efficiently transferred between cells, as a conjugative plasmid cargo, via conjugation. We first tested the conjugative transfer of the pRK24 plasmid carrying the DGR/VR cassette into a recipient strain with spectinomycin resistance. By using a spotting assay with antibiotic selections, we observed that the modified pRK24 plasmid could transfer into the recipient strain with a conjugation rate of 50-100% as quantified relative to the donor **(Fig. 3B, S12)**, consistent with the observations in retron recombineering (*24*). This high level of conjugative transfer is desirable to ensure that library diversity is not bottlenecked by conjugative transfer. Next, we tested whether the DGR system components could function with the template RNA transcribed from the TR in the recipient cell after conjugative transfer to drive DGR-mediated mutagenesis of the sfGFP (VR) on the conjugative plasmid. Indeed, we observed DGR component-dependent mutations within the targeted region of the *sfgfp* gene at expected levels, demonstrating that our DGR system is fully compatible with conjugation of DGR components and VR into the TR-bearing recipient strain **(Fig. 3C)**. Finally, we sought to identify the best antibiotic resistance markers for the TR vectors and optimize the induction of CspRecT/mMutL expression. We found that the plasmid harbouring nourseothricin (NTC) resistance led to the most VR mutations in the context of the conjugative plasmid carrying the DGR system **(Fig. S13)**. We found that CspRecT and mMutL led to cell death during DGR induction with 0.42-6.67 mM arabinose **(Fig. S14A)**. However, a low level of induction (0.42 mM) for 10 hours was sufficient for DGR-mediated mutagenesis with minimal cell death **(Fig. S14B)**. We also found that the TR vector affected the survival rate after DGR induction. Among all three plasmids with antibiotic resistances we tested, again the NTC-resistant plasmid led to the least cell death **(Fig. S14C)**.

Next, we designed a sustainable DGR-mediated hypermutation strategy based on iterating conjugation steps with interspersed induction of the DGR system **(Fig. S15A)**. To achieve this HGT-DGR cycle, we need to enrich the recipients after each conjugation step by alternating antibiotic selection. We prepared the TR host strains by inserting three different resistance genes (*cat* for chloramphenicol, *aadA* for spectinomycin, *hph* for hygromycin) into the *E. coli* genome. Next, we tested the conjugation efficiencies for the pRK24 harbouring sfGFP and the constant DGR components into these strains which revealed that the chloramphenicol and spectinomycin-resistant hosts led to the highest conjugation efficiencies **(Fig. S15B)**. By iteratively conjugating conjugative plasmids carrying the *sfgfp* gene into a recipient carrying the corresponding TR and alternating antibiotic markers to avoid cross-contamination with donor cells, we observed the accumulation of mutations in the adenine positions of the VR at the population level, and the replenishment of the TR **(Fig. 3D, S15C)**. After 7 days of continuous conjugation and DGR induction, the mean mutation frequency at all adenine positions within the VR reached >6% **(Fig. 3D)**.

To determine the precise mutation distribution across the population (i.e. to determine whether mutations were concentrated in a small set of highly mutated variants or distributed across a broader and more diverse mutant population), we performed a clonal sequencing experiment on 86 single colonies after 7 days of conjugation and DGR induction. The resulting per-adenine mutation frequency spectrum was consistent with the global average NGS data **(Fig. S15D)**, with up to 40% of clones (34 / 86) carrying VR mutations at adenine positions **(Fig. 3E)**. Most variants contained 1–4 mutations **(Fig. 3F)**, and pairwise Hamming distances among variant fingerprints—strings obtained by concatenating the nucleotide calls (A/C/G/T) at each site in the observed variable-site panel —were broadly distributed (median 4; IQR 3–5) **(Fig. 3G)**, consistent with dispersed combinatorial diversity and only rare duplication.

### Evolution of aminoacyl-tRNA synthetase (aaRS) via HGT-DGR

To demonstrate the utility of our HGT-DGR system for targeted hypermutation, we sought to evolve the active site of pyrrolysyl-tRNA synthetase from *Methanococcus mazei* (*Mm*PylRS). *Mm*PylRS is the most widely used aminoacyl-tRNA synthetase (aaRS) for genetic code expansion (*42*), and we challenged it to specifically charge two non-canonical amino acids (ncAAs): Nε-benzyloxycarbonyl-L-lysine (CbzK) and Nε-benzyl-L-lysine (BenzK), as also described for retron editors (*24*).

We introduced the *Mm*PylRS gene (as the VR) on the conjugative plasmid bearing the constitutive DGR components, and designed TR sequences targeting defined codons in the PylRS active site. First, we conjugated the conjugative plasmid into recipient cells bearing the TRs and induced DGR mutagenesis to generate a diverse library of *Mm*PylRS active-stie mutants. Next, we conjugated this library into a strain containing the sfGFP-cat reporter system for selection and selected for cells that were both chloramphenicol resistant and exhibited green fluorescence **(Fig. 4A)**.

**Figure 4.**
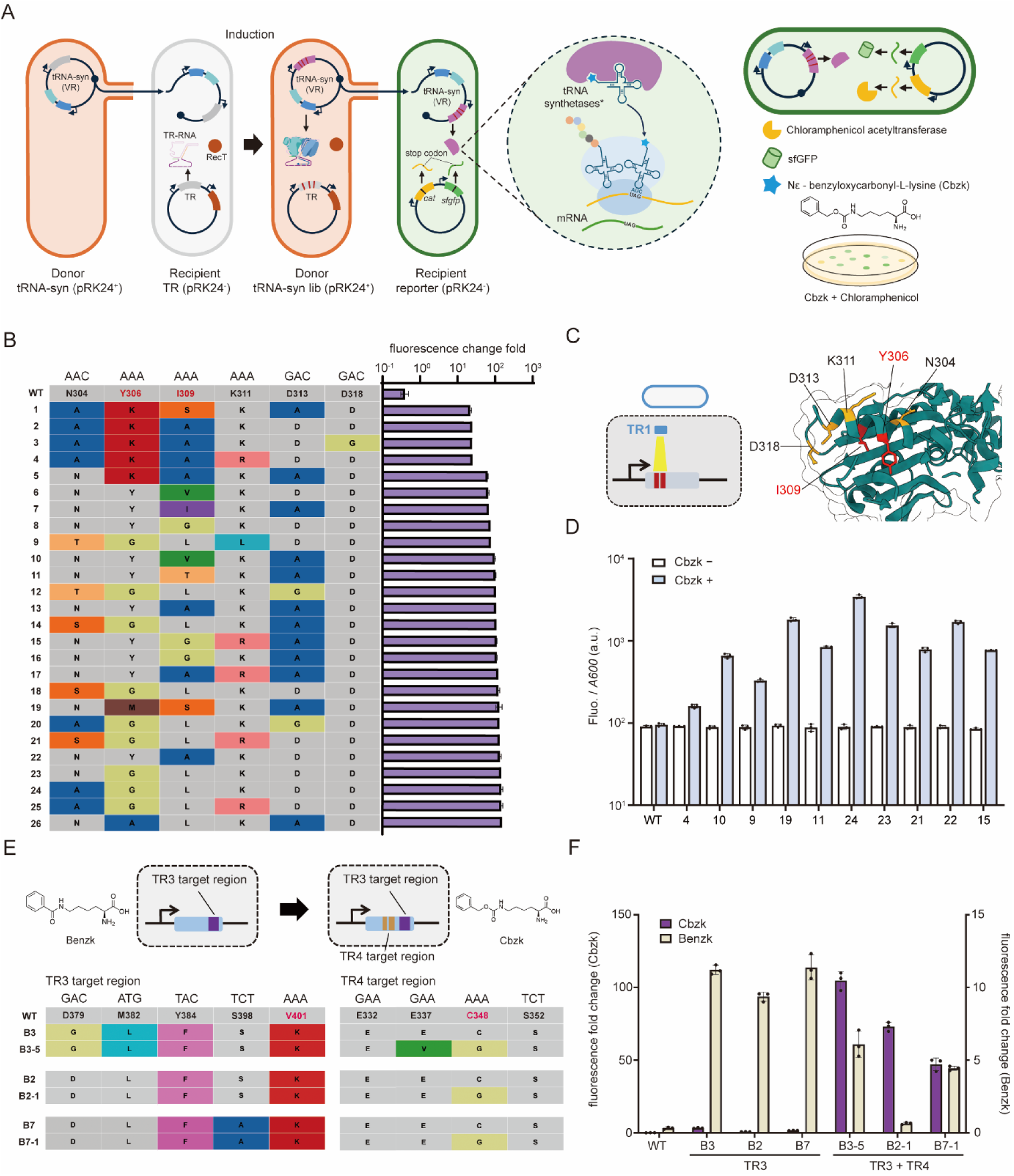
Evolution of Mm PylRS for incorporation of non-canonical amino acid via HGT-DGR. **A,** Schematic of aaRS evolution and the reporting system with a dual GFP/chloramphenicol-resistance reporter. **B,** Table of amino acid mutations enriched from a CbzK evolution experiment with the dynamic range of CbzK dependent expression of GFP shown to the right of the corresponding sequence. **C,** Schematic of TR1 targeting the Mm PylRS gene. The intentionally targeted codon is shown in red while the non-targeted codon in yellow mapped in the structure of Mm PylRS predicted by AlphaFold. **D,** Characterization of CbzK-dependent expression using the 10 different variants shown in **B**. **E,** Table of mutations enriched from the BenzK to CbzK evolution experiment. The labelling before dash indicates the ancestor of the clone screened using BenzK, while the number after dash indicates the corresponding progeny to the enriched clones. **F,** Fluorescence fold change for CbzK and BenzK responsive clones from the TR3 evolution and TR4 evolution. Error bars, s.d. (n = 3)

As a first proof of concept, we sought to evolve *Mm*PylRS for the ncAA substrate Nε-benzyloxycarbonyl-L-lysine (CbzK). We designed a TR containing an AAA triplet at each position corresponding to the codons for residues Y306 and I309 (TR1, covering *Mm*PylRS residues F295–F323), as the Y306G mutant of *Mm*PylRS is known to accommodate CbzK (*43*). After just one round of evolution, we enriched several hits with CbzK-dependent GFP expression. Among the 54 clones sequenced, we identified 26 unique variants,16 of which showed variation at position 306. Among these 9 out of 16 showed the Y306G mutation, consistent with previous results, and 5 out of 16 showed Y306K (encoded by the AAA codon, derived from the TR without mutation). All 26 clones contained at least one non-AAA mutation in positions other than 306 and 309. Interestingly, we also enriched some mutations outside codons 306 and 309 in PylRS; these changes likely result from the mutation of other adenines in the VR **(Fig. 4B, C)**, which was expected as all adenines in the TR are subject to mutagenesis. We also lowered the concentration of CbzK (0.2 mM) to have more stringent ncAA-dependent GFP readouts. All 10 representative clones tested showed significant ncAA-dependent GFP expression, confirming the robustness of our selection method **(Fig. 4D)**.

Conjugation of the gene of interest containing conjugative plasmid into recipients harbouring TRs conveniently allows the use of distinct TRs in each iteration. As a proof of concept, we selected two regions of the *Mm*PylRS gene as targets and designed corresponding TRs (TR3 and TR4, covering *Mm*PylRS aa V377-R409 (TR3) and G326-R356 (TR4) respectively). We sequentially transferred the conjugative plasmid carrying the PylRS gene into recipients carrying TR3 and TR4. After each conjugation step, we induced DGR mutagenesis and observed that DGR derived mutations arose successively on the two VRs in the population **(Fig. S16)**.

Next, we decided to evolve PylRS to accept a more challenging substrate, N^ε^-benzyl-L-lysine (BenzK). On the basis of previous work on BenzK-selective synthetases, we designed two different TRs, targeting C348 (TR2, covering *Mm*PylRS aa R330-L359) and V401 (TR3) with an AAT and AAA respectively. We mixed the TR2 and TR3 containing cells as recipients during conjugation **(Fig. S17A)**. After 2 rounds of conjugation and DGR mutagenesis (HGT-DGR), followed by selection, we were able to enrich 3 unique solutions. Then we ran three additional iterations of HGT-DGR with the total of 5 iterations followed by selection. This yielded 4 additional PylRS variants that enable the incorporation of BenzK. All enriched clones had mutations within the TR3 region but did not have mutations within the TR2 region **(Fig. S17B)**.

We then transferred the small library of clones we discovered for BenzK-incorporation, with mutations in TR3, into a recipient strain harboring TR4, which targets M344 and C348 with AAA and AAC respectively. Then we conducted the DGR mutagenesis and conjugated into the recipient with the reporter for selection of CbzK-charging aaRS variants **(Fig. S18A)**. We discovered that mutations in TR4 target regions, when combined with the previously discovered TR3 mutations, conferred CbzK-responsive aaRS activity **(Fig. 4E, F, S18)**. Some of the variants we obtained had a shift in substrate preference from BenzK to CbzK, while others retained more than 50% activity for BenzK, and showed a broadened substrate preference **(Fig. 4F, S19)**. We selected B3 and B3-5 for further mass spectrometry analysis, which confirmed the ability of B3-5 to utilize both Cbzk and Benzk **(Fig. S20)**.

## Discussion

DGRs are widespread in bacteria and viruses, particularly in environments such as the animal gut, and are capable of a level of hypermutation that exceeds that of the vertebrate immune system. Here, using a synthetic retrohoming strategy, we show that DGR-mediated hypermutation can be realized in *E. coli* for genes of interest as also described (*19*). We find that DGR does not just enable the targeting of mutations to a specific gene segment (as specified by the TR), but to specific A nucleotides within that segment. This not only enables the precise targeting of mutations, but opens up the possibility to *a priori* tune different protein regions for higher or lower mutability simply by recoding with a low or high A-content (e.g. encoding Arg as AGA rather than CGU etc.) or for different mutational outcomes.

Our results demonstrate that, in cases targeting a few defined codons, the mutation rate mediated by a single round of DGR may be sufficient to support directed evolution. However, to cover a larger gene segment and explore a wider range of sequence variants, a higher mutation frequency is needed. This is not possible with the standard synthetic DGR system as installed in *E. coli* **(Fig. 1)**, as it is unstable under prolonged continuous culture and mutations in the target VR sequence cease to accumulate after ca. 3 days. We show that horizontal gene transfer using conjugation (HGT-DGR) is a strategy to address these problems by continuously transferring target sequences to refresh the host background and TR. This allows the mutant sequences accumulated to comprise 40% of the population within 7 days or >10^9^ variants that are generated in one week in just 1 mL of bacterial culture. Integrating HGT-mediated transfer of DGR-mutated libraries enables continuous mutation at the same or multiple user-defined loci within the population, which enables rapid biomolecular engineering as we demonstrate by the directed evolution of the *Mm*PylRS aaRS to charge new ncAA substrates.

A future focus will be on continuing to increase the efficiency of DGR-based mutagenesis to enable more rapid mutation of multiple TRs in single clones. Optimal mutation rates are likely target and case-specific and HGT-DGR allows these to be tuned both by rapid, iterative choice of TR encoding and host-strain. Current mutation rates of HGT-DGR are modest (per round) and do not cover 100% of the cell population. However, we would argue that this is advantageous as, in conjunction with selection, too high levels of mutation are generally undesirable as they may lead to large numbers of non-functional variants as well as to an overwriting of the existing enriched mutations in VRs rather than increasing overall diversity. Finally, the maintenance of a significant number of wild-type (wt) sequences in the population may be advantageous as backcrossing with wt (through recombination) has been shown to accelerate adaptive evolution by enhancing the spread of beneficial mutations and mitigating phenotypic degradation through the accumulation of mildly deleterious mutations (mutational drift) (*44*).

We expect that our DGR-based continuous evolution system will be a valuable addition to the toolbox available for directed evolution experiments, enabling continuous mutagenesis over discrete user-defined sequence windows. We note that our system may be particularly suitable for the rapid evolution of binding interfaces (or catalytic sites) specified by discrete sequence segments such as the CDR loops in antibody-antigen and TCR-MHC binding interfaces. Future work will focus on further developing the mutagenic efficiency of the system and on automating cycles of HGT and DGR-mediated hypermutation with the goal of enabling massively parallel continuous evolution experiments.

A particularly exciting opportunity arising from HGT-DGR is its potential to support the generation of training data for artificial intelligence (AI) models. By progressively increasing the proportion of mutated individuals relative to the wild type in the population, while restricting diversification to precisely defined sequence windows, HGT-DGR can generate large libraries with well-defined mutational spaces. When coupled to high-throughput functional characterization, this provides a distinctive approach for the automated generation of highly structured sequence–function datasets for training and benchmarking AI models. Future work will therefore explore the use of HGT-DGR to substantially reduce the experimental cost of producing large-scale, information-rich datasets for AI-guided biomolecular engineering.

## Acknowledgments

The authors thank Dr. Kelly Nguyen for assistance with the manuscript preparation. The authors thank Dr. Kim C Liu for providing NGS-related technical support. The authors also thank the Mass Spectrometry facility, Flow Cytometry facility and the Media Prep and Glasswash facility of MRC-LMB for their technical support.

For the purposes of open access, the MRC Laboratory of Molecular Biology has applied a CC BY public copyright license to any Author Accepted Manuscript version arising from this work.

## Funding

Medical Research Council (MRC) program MC_U105178804 (PH)

Medical Research Council (MRC) program MC_U105181009 and MC_UP_A024_1008 (JWC)

Wellcome Trust Investigator Award 220808/Z/20/Z (JWC)

MRC–AstraZeneca Blue Sky Grant (YL, GA)

European Molecular Biology Organization ALTF 93-2023 (YG)

National Health and Medical Research Council (NHMRC) Australia Investigator Grant GNT2018461 (F.R)

The funders had no role in study design, data collection and analysis, decision to publish or preparation of the manuscript.

## Author contributions

Conceptualization: YL, YG, PH

Methodology: YL, YG, FR, MS

Investigation: YL, YG, FR, GA

Funding acquisition: PH, JWC

Supervision: PH, JWC

Writing - original draft: YL, YG, FR

Writing - review & editing: YL, YG, GA, FR, MS, PH, JWC

## Competing interests

YL was supported by the MRC–AstraZeneca Blue Sky Grant. AstraZeneca reviewed the manuscript prior to submission but did not provide input or request changes. The authors declare no other competing interests.

## Data and materials availability

All the authors are listed as inventors on patent applications submitted by MRC-LMB. Plasmids and strains listed in tables S1 and S2 are available from the authors upon reasonable request.

## Supplementary Materials

Materials and Methods

Figs. S1 to S22

Tables S1 to S4

References (45–48)

## Notes

### Summary of Updates

Updated terminology describing the conjugation system to accurately reflect the use of the pRK24 conjugative plasmid rather than the F plasmid throughout the manuscript and supplementary materials. Corrected the spelling of author Jason Chin's name.

