## Supplementary material for "Continuous target-specific mutagenesis and rapid gene evolution by diversity-generating retroelements in *Escherichia coli*": Sup_info

Yang Liu *et al.*

**The PDF file includes:**

Materials and Methods

Figs. S1 to S22

Tables S1 to S4

References (45-48)

### Materials and methods

#### Construction and strains

The construction of all plasmids except the conjugative-plasmid and genome editing were performed using conventional molecular biology methods. The *avd*, *brt*, and *mtd* genes, TR, and the sequence of HDV ribozyme were synthesized by commercial company (IDT). The DGR generator pLG-2 was constructed by synthesized *avd* and *brt* gene, the $P_{BAD}$ promoter amplified from pLY124 (*45*), and the vector is from pLY152 (*45*). For the DGR generator pLG-1, it was constructed by replacing the $P_{BAD}$ promoter of pLG-2 to $P_{rhaB}$ promoter which was amplified from pLY151 (*45*). The pLG6 - pLG9 was assembled by vector, *mtd* gene (WT VR), $P_{lux}2$ promoter, WT TR, and HDV ribozyme. The $P_{lux}2$ promoter of pLG6 - pLG9 is amplified from pLY47 (*45*). The vector of pLG6 - pLG9 is from pLY151. For pLG-10 - pLG-12, the DGR cassette was amplified from pLG-1. The *sfgfp* gene is from pLY152, the vector is from pLG-6. All the plasmids are listed in the **Table S1**. The structures of all the plasmids have been summarized in **Fig. S21** and **Fig. S22**.

NEB 10-beta *E. coli* was used for plasmid construction. A DH10B *E. coli* strain with Δ*sbcB*, Δ*recJ*, Δ*exoX* genotype, and with different antibiotics resistance was employed for the HGT-DGR experiments. All the *E. coli* strains have been listed in **Table S2**.

All the TR and VR sequences used in this study have been listed in **Table S4**.

#### Conjugative-plasmid (modified pRK24) construction

Conjugative-plasmid used in this study was derived from an RK2-derived conjugative plasmid pRK24 (*46*) where a PH (PheS/Hygromycin) marker was strategically inserted into a specific region on the pRK24. Then the VR and DGR were introduced into the same region to replace the PH marker via a lambda red recombination approach. In brief, a modified pEcCas9 plasmid (*47*) with chloramphenicol resistances was transformed into the strain with pRK24. Then the cells with both plasmids were inoculated into an overnight culture with hygromycin (100 μg/mL) and chloramphenicol (20 μg/mL). The overnight culture was then inoculated 1:50 into 100 mL of fresh 2XYT media with apramycin (50 μg/mL) and chloramphenicol (20 μg/mL) until OD ~0.3. Then 10 mM final concentration of arabinose was added to the culture to induce the lambda red proteins. After 30 minutes induction, the cells were harvested and prepared cold into electrocompetent cells. For each 100 μL competent cells, ~2 μg PCR products containing VR, DGR and Kan resistance genes were introduced into the cells via electroporation. The cells were recovered in 5 mL SOB for 3 hours before plating on the 2XYT agar selection plate with Kanamycin resistance (50 μg/mL) and 4-chlorophenylalanine (25 mM). Then the colonies grown on the selection plate were picked and respotted onto a fresh 2XYT agar plate with the same selection. Then each newly grown spots were picked for PCR and nanopore sequencing to confirm the correct insertion of DGR and VR on the pRK24. Then the correct clones were restreaked on a fresh 2XYT agar plate with Kan and 7.5% sucrose to obtain single colony.

#### WT and GFP DGR tests:

Three single colonies were inoculated into 200 μL overnight culture in LB with appropriate antibiotics, ampicillin (100 μg/mL), kanamycin (50 μg/mL), hygromycin (100 μg/mL), within a flat bottom clear 96-well plate (Starlab). The overnight culture was then re-inoculated into the same 200 µL of fresh LB media with the antibiotics at 1 to 100 dilutions. The culture was prepared with or without inducers. The inducers were added with the following final concentrations unless specified otherwise: L-arabinose (0.1%), L-rhamnose (10 mM), N-(3-Oxohexanoyl)-L-homoserine lactone, AHL (12.5 µM).

Then the culture was incubated in a table top shaker at 37°C, 800 rpm for 24 hours.

**GFP evolution on HGT-DGR:**

Three single colonies were inoculated into 200 µL overnight culture in LB with antibiotics: kanamycin (50 µg/mL) and nourseothricin (100 µg/mL) within a flat bottom clear 96-well plate (Starlab). The overnight culture was then re-inoculated into the same 200 µL of fresh LB media with the antibiotics at 1 to 100 dilutions. The culture was prepared with or without inducers. The inducers were added with the following final concentrations unless specified otherwise: L-arabinose (0.1%), L-rhamnose (10 mM). Then the culture was incubated in a table top shaker at 37°C, 800 rpm for 24 hours.

Then cells were collected for downstream NGS or spotting assay.

For the 7-days continuous evolution of GPF, the conjugative-plasmid pLG-13 was first conjugated into a recipient *E. coli* strain (DH10B) carrying TR generator plasmid pLG-17. This transconjugant was streaked to obtain single colonies.

For the conjugation group, three single colonies were picked into 20 mL 2XTY media with nourseothricin (100 µg/mL) and inducers (L-arabinose (0.00625%), L-rhamnose (20 mM)) in a 200 mL flask for 10 hours culture (37°C, 180 rpm). Meanwhile, 3-4 hours before the end of induction, inoculate overnight cultured recipient cells (DH10B with ΔsbcB, ΔrecJ, ΔexoX genotype with spectinomycin resistance) at a ratio of 1:100 into 5 mL 2XTY media with spectinomycin (50 µg/mL) in a 50 mL Facon tube, and incubate at 37°C, 180 rpm. After the induction, collect the induced donor cells and recipient cells (equivalent to 1 mL of OD = 1) into 1.5 mL EP tube, separately. Then, perform the standard conjugation. In subsequent experiments, 10 hours induction with overnight enriched transconjugant was repeated daily, alternating conjugations into recipient with chloramphenicol or spectinomycin resistance.

For the subculture group, three single colonies of the first transconjugant were picked into 20 mL 2XTY media with nourseothricin (100 µg/mL) and inducers (L-arabinose (0.00625%), L-rhamnose (20 mM)) in a 200 mL flask for 10 hours culture (37°C, 180 rpm). After the induction, collect the induced cells (equivalent to 1 mL of OD = 1) into 1.5 mL EP tube. Centrifuge at 6000 rpm for 3 min, discard the supernatant and resuspend the cells in 1 mL of PBS. Centrifuge at 6000 rpm for 3 min, resuspend the cells in 200 µL of 2XTY media, and inoculate into 20 mL of 2XTY media with kanamycin (50 µg/mL) and nourseothricin (100 µg/mL) for overnight culture. In subsequent experiments, 10 hours induction and overnight culture was repeated daily. 10 µL of culture were collected from each flask daily before and after induction, for downstream NGS or spotting assay.

**Conjugation and spotting assay**

The donor and the recipient cells in liquid culture were normalized into equal amounts of cells equivalent to 0.5 mL of OD=1 cell culture. Both donor and recipient were first washed with PBS twice and then mix and wash with PBS one more time. After spinning down the donor recipient mixture, the cells were resuspended in minimal amount of PBS and dropcast onto a pre-cut cellulose filter paper (company) with 2 XYT agar media underneath. Then the cell aggregate was air dried until no droplet of liquid was seen and then was incubated at 37 C for 1 hour for conjugation. Then the cells were resuspended from the filter paper and vortex to break the mating pair. Then the resuspended cells recovered in 5 mL SOB for an hour before spotting on the agar plate with different antibiotics selection.

To quantify the survival rate, the cells were first subject to serial 5X dilution. And then 5 μL of the undiluted and diluted cell culture were directly spotted onto the agar plate with corresponding antibiotics. The survival rate was calculated by dividing the colony numbers from the group with inducers to the group without inducers.

To quantify conjugation efficiency, the cells were first subjected to a series of 5X dilutions. And then 5 μL of the undiluted and diluted cell culture were directly spotted onto agar plates containing different antibiotics (antibiotics for the conjugative-plasmid, antibiotics for the TR plasmid, and antibiotics for the transconjugant). The true donor count can be obtained by subtracting the number of transconjugants from the total number of cells carrying the pRK24 antibiotic resistance marker. Dividing the number of transconjugants by the true donor count yields the conjugation efficiency over the donor. Dividing the number of transconjugants by the recipient count yields the conjugation efficiency over the recipient.

**NGS and mutation analysis**

For each bacterial sample, 10 μL of cells were spun down and resuspended in 10 μL of QuickExtract™ DNA Extraction Solution (LGC Biosearch Technologies) and run through 65 C 10 mins followed by 98 C 5 mins in the thermo cycler. Then 1 μL of extracted sample was used as a PCR template with Q5 polymerase for amplification of VR and TR with their corresponding primers. Then the PCR samples was subject to 1 hour of DpnI digestion at 37C. The PCR sample was then purified and library prepared for using for NGS sequencing. All the NGS was running on a NextSeq illumnia sequencing machine with either P1 100 or P2 100 cartridge at the core facility at LMB. All the primers for NGS sample preparation have been listed in **Table S3**.

The raw NGS reads were first trimmed off their adapters using cutadapt and then the trimmed reads were aligned using bowtie2 with an in-house script. The aligned reads were further analyzed using an in-house python script to extract the mutation bases and rate for each aligned mutation. All the data were then combined, and the figure was rendered in prism.

**CbzK and BzK evolution**

To evolve the tRNA synthetase, the conjugative-plasmid containing tRNA synthetases was first conjugated into a TR containing triple KO strain. The cells harbouring both plasmids were inoculated in an overnight culture containing conjugative-plasmid and then re-inoculated into a 20 mL of 2XYT culture containing 20 mM rhamnose, 0.00625% of arabinose and 100 μg/ml of nourseothricin and induced for 10 hours. For multiple rounds of evolution, 1 ml OD =1 equivalent of induced cells were immediately prepared as a donor to conjugate to the same amount of recipient cells containing TR plasmids. After conjugation and recovery, the cells were inoculated into 20 mL of fresh media with Kanamycin (50 μg/mL), nourseothricin (100 μg/ml), and appropriate antibiotics for the recipient (50 μg/mL Spectinomycin/25 μg/mL Chloramphenicol/none) to enrich the transconjugant overnight. In the next day, the overnight culture was diluted (1:100 dilution) into 20 mL 2XTY with nourseothricin (100 µg/ml), appropriate antibiotics for the recipient (50 µg/mL Spectinomycin/25 µg/mL Chloramphenicol/none), 20 mM rhamnose, and 0.00625% of arabinose for 10 hours induction.

For the selection, 1 ml OD =1 equivalent of induced cells were immediately prepared as a donor to conjugate to the same amount of recipient cells containing reporter plasmids. After conjugation and recovery, the cells were inoculated into 20 mL of fresh media with tetracycline (10 µg/mL) and Kanamycin (50 µg/mL) to enrich the transconjugant.

200 µL of overnight culture was then reinoculated into 20 mL of fresh 2XYT culture containing 2 mM CbzK/BzK, 10 mM arabinose, 50 µg/ml Kanamycin, 10 µg/mL tetracycline and induce for 6 hours at 37 °C with shaking at 180 rpm. After induction, the cells were plated onto the 2XYT selection agar plate with 2 mM CbzK/BzK, 10 mM arabinose, 50 µg/ml Kanamycin, 10 µg/mL tetracycline and 70 µg/mL of chloramphenicol.

GFP positive colonies were then picked for initial screening and further characterization. For initial screening, we inoculated the single colonies into 200 µL of 2XTY medium with 50 µg/ml Kanamycin, 10 µg/mL tetracycline for overnight culture. The overnight culture was diluted 100-fold into 200 µL 2XTY medium in 96-well plate with 50 µg/ml Kanamycin, 10 µg/mL tetracycline, arabinose (10 mM), with or without 2 mM non-natural amino acids (Cbzk/Benzk), for a 5.5-hour culture. Then the cells were resuspended in an equal volume of PBS in 96-well plate. We read the plate for OD (*A600*) and fluorescence intensity (FI) (Excitation 485 nm/Emission 520 nm, gain 200) by plate reader.

**Proof of TR reprogramming concept**

The conjugative-plasmid pLG-14 was first conjugated into a recipient *E. coli* strain (DH10B with Δ*sbcB*, Δ*recJ*, Δ*exoX* genotype) carrying plasmid pLG-20. The overnight cultured transconjugant was 1:100 diluted into 20 mL 2XTY media with nourseothricin (100 µg/mL) and inducers (L-arabinose (0.00625%), L-rhamnose (20 mM) in a 200 mL flask for 10 hours culture (37°C, 180 rpm). Meanwhile, a single colony of a recipient *E. coli* strain (DH10B with ΔsbcB, ΔrecJ, ΔexoX genotype with spectinomycin resistance) carrying plasmid pLG-21 was picked into 5 mL 2XTY media with spectinomycin (50 µg/mL) and nourseothricin (100 µg/mL) in a 50 mL Facon tube, and incubate at 37°C, 180 rpm. After the induction, collect the induced donor cells and recipient cells (equivalent to 1 mL of OD = 1) into 1.5 mL EP tube, separately. Then, perform the standard conjugation. 10 µL of culture were collected from each flask daily before and after induction, for downstream NGS or spotting assay.

**CbzK/Bzk synthetase characterization**

The positive colonies (with green color) were picked and inoculated into overnight 2XYT culture with tetracycline and kanamycin. The overnight culture was then re-inoculated (1:100 dilution) into 200 µL fresh media containing tetracycline, kanamycin, and 10 mM arabinose and with or without ncAA (0.2 mM Cbzk for TR1 evolution and 2 mM Cbzk/Benzk for TR3 and TR4 evolution). After 5.5 hours of induction, the cells were collected, resuspended in PBS and measured the GFP readout (Gain = 100 for TR1 evolution experiment and Gain = 200 for TR3 and TR4 evolution experiments). The fluorescence fold change was calculated based on the fold change of GFP with/without ncAA. The calculation method is to divide the difference between the single-cell fluorescence under nnAA-positive and the single-cell fluorescence under nnAA-negative by the single-cell fluorescence under nnAA-negative conditions.

**Generative AI use**

The authors used ChatGPT (OpenAI) for limited assistance with drafting and debugging portions of the analysis code and for copy editing of author-generated text. All AI-assisted code and text were independently reviewed, tested and revised by the authors, who take full responsibility for the final manuscript, code and analyses.

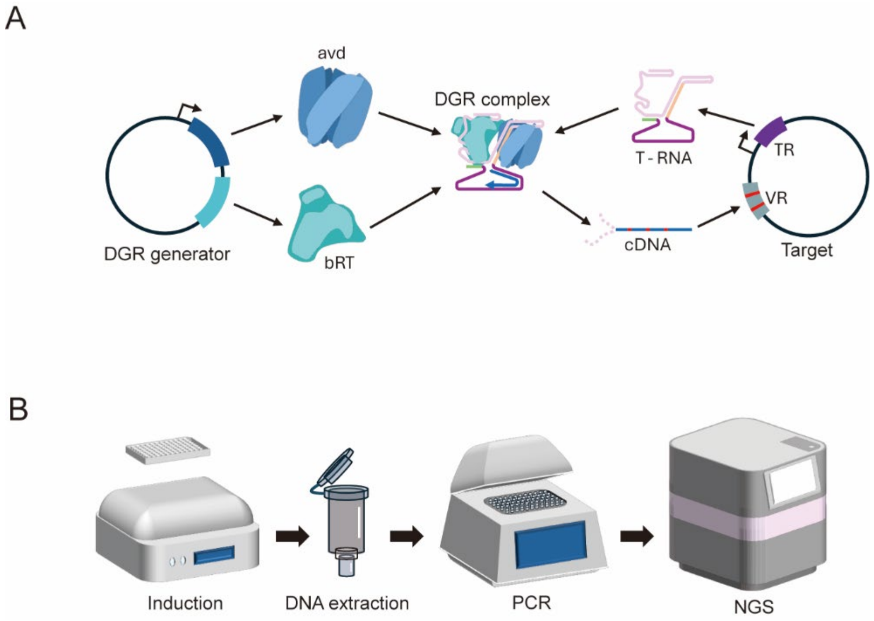


**Figure S1. Schematic of the design and characterization workflow of the DGR system.** **A,** Schematic of two-plasmid system for DGR function *in vivo*. **B,** Schematic of NGS workflow.

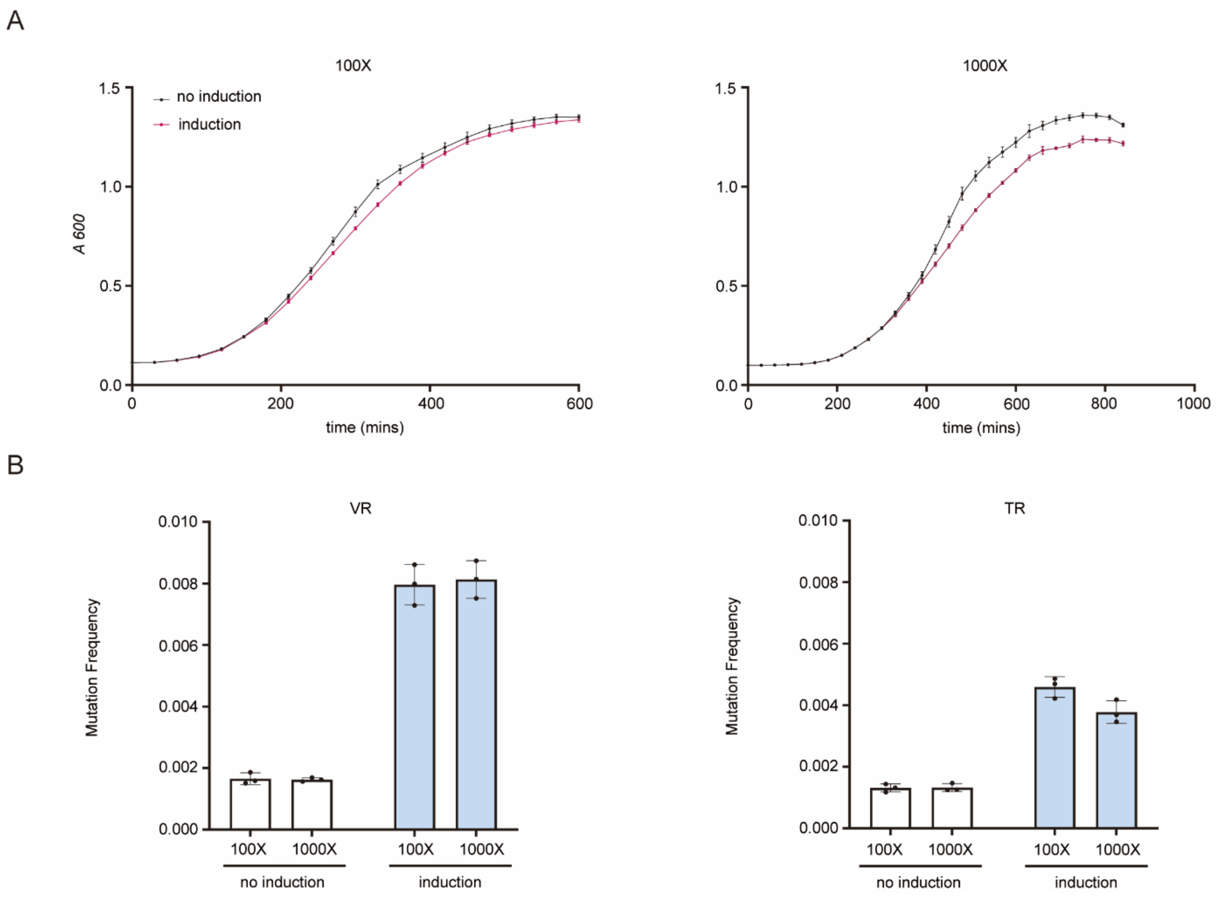


**Figure S2. Cell generation does not affect the frequency of a single DGR induction process. A,** Cell growth curves under single DGR induction. The initial cell concentrations were 100-fold dilution overnight culture (left) and 1000-fold dilution overnight culture (right). **B,** Mutation frequency of adenine sites on VR (adenine sites present only in both TR and VR, left) or TR (right). The final OD values (at sampling point) were all between 1.3 and 1.4, and the sampling times were 10h (for 100X) and 14h (for 1000X) to ensure that the final OD values were consistent. Error bars, s.d. ($n = 3$)

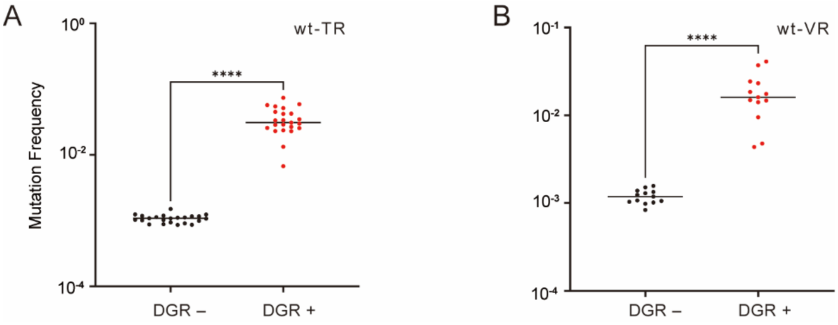


**Figure S3. Mutation frequency (24h) of adenine sites on TR and VR. A,** Mutation frequency of adenine sites on TR. **B,** Mutation frequency of adenine sites on VR (adenine sites present only in both TR and VR). Statistical difference was determined by a two-tailed Welch's *t* test for TR, $p < 0.0001$, $t = 10.41$, and two-tailed Welch's *t* test for VR, $p < 0.0001$, $t = 5.699$. *p* value summary: ****$p$ value $< 0.0001$, $0.0001 <$ ***$p$ value $< 0.001$, $0.001 <$ **$p$ value $< 0.01$, $0.01 <$ *$p$ value $< 0.05$, $p$ value $\geq 0.05$: n.s. Error bars, s.d. ($n = 3$)

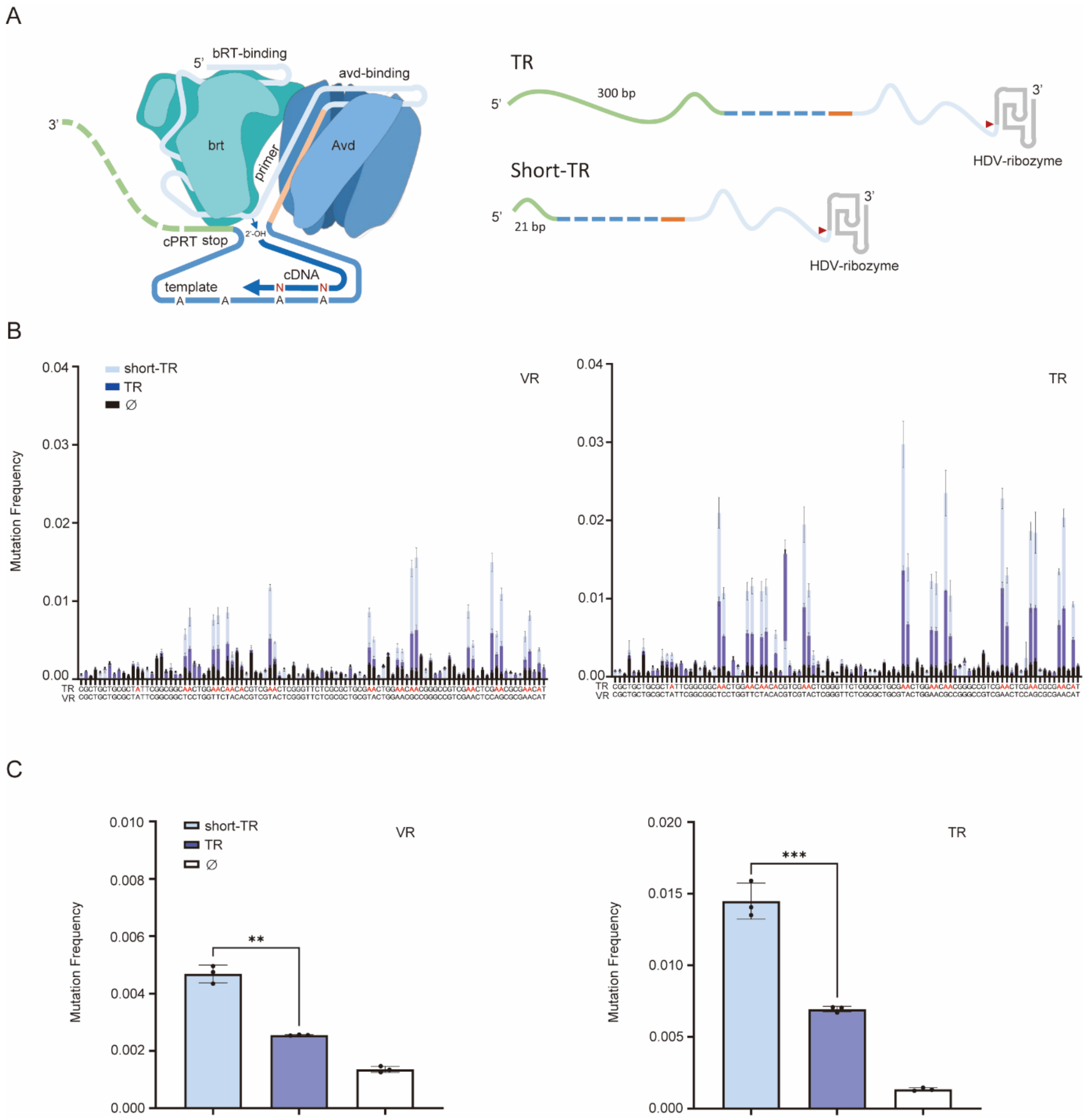


**Figure S4. Short TR 5' sequence is more efficient for DGR retrohoming. A,** Two different designs of TR maintaining 300 bp (TR) or 21 bp (short-TR) 5' upstream from the natural BBP1 TR sequence. **B,** Mutation frequency (5h) barplot for short-TR, 300 bp TR (TR) and no DGR cassette control (∅). **C,** Mutation frequencies (5h) calculated from the average of all A position mutations from VR and TR sequences. Statistical difference was determined by a two-tailed Welch's *t* test for VR, $p = 0.0068$, $t = 11.97$, and two-tailed unpaired *t* test for TR, $p = 0.0005$, $t = 10.27$. *p* value summary: $****p$ value $< 0.0001$, $0.0001 < ***p$ value $< 0.001$, $0.001 < **p$ value $< 0.01$, $0.01 < *p$ value $< 0.05$, $p$ value $\geq 0.05$: n.s. Error bars, s.d. ($n = 3$)

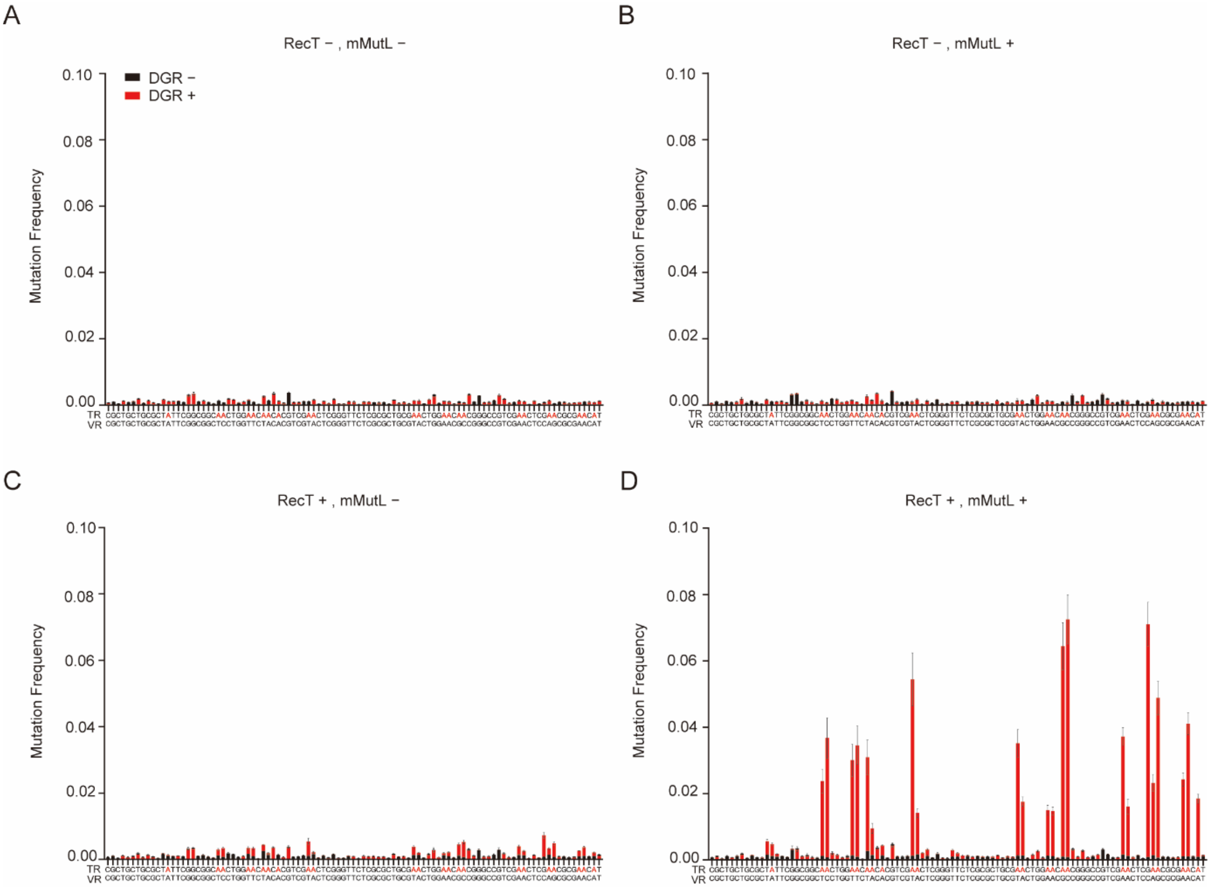


**Figure S5. Both mMutL and CspRecT are essential for DGR-mediated mutations.** Mutation frequency (24h) barplots of VR with induction (DGR +) or no induction (DGR -) using the natural BBP1 VR and TR sequences, **A,** without mMutL (mMutL-) and CspRecT (RecT-). **B,** with mMutL (mMutL+) and without CspRecT (RecT-). **C,** without mMutL (mMutL-) and with CspRecT (RecT+). **D,** with both mMutL (mMutL+) and CspRecT (RecT+). Adenine positions present in the TR are highlighted in red in the sequences below the barplots. This data is also shown in **Fig. 1B.** Error bars, s.d. ($n = 3$)

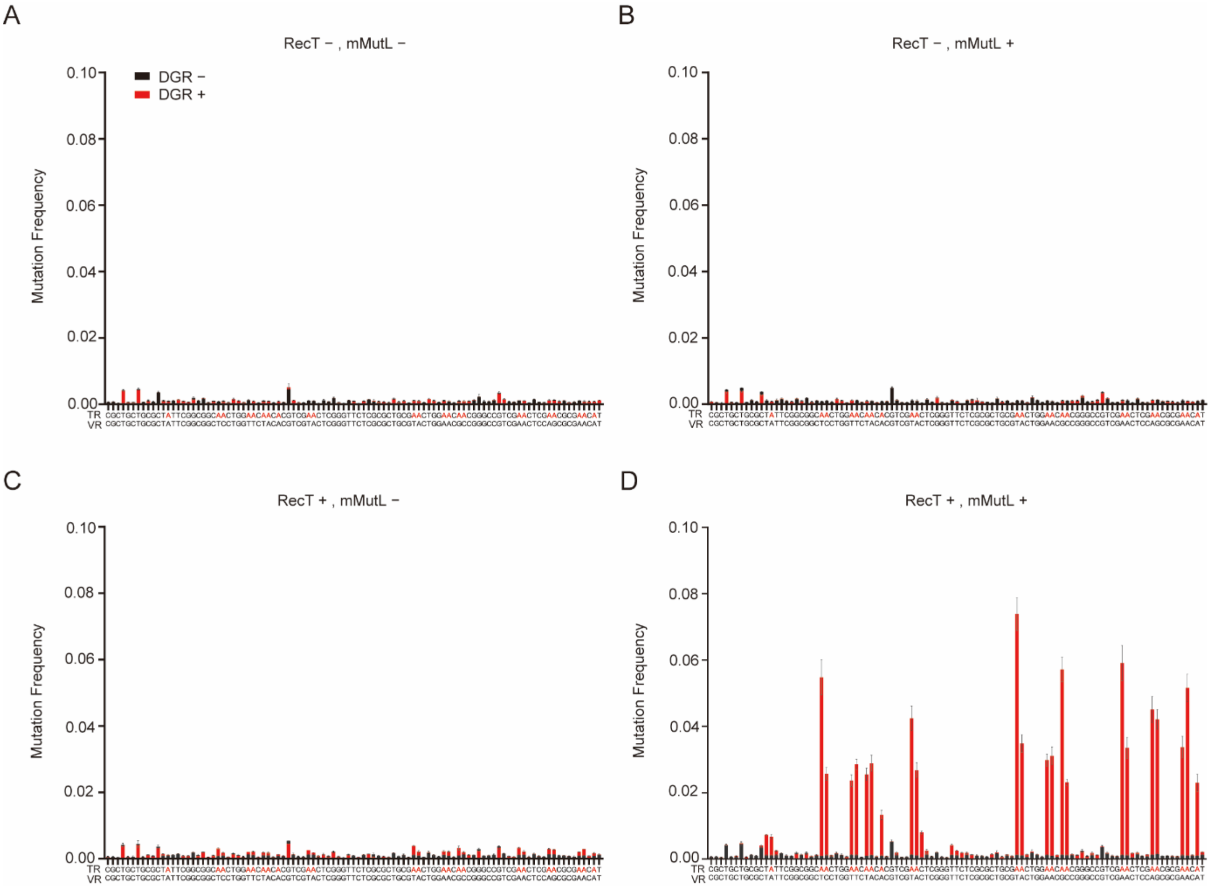


**Figure S6. Mutation Frequency (24h) Barplot of TR with induction (DGR +) or no induction (DGR-) using the natural BBP1 VR and TR sequences**, **A,** without mMutL (mMutL-) and CspRecT (RecT-). **B,** with mMutL (mMutL+) and without CspRecT (RecT-). **C,** without mMutL (mMutL-) and with CspRecT (RecT+). **D,** with both mMutL (mMutL+) and CspRecT (RecT+). Adenine positions present in the TR are highlighted in red in the sequences below the barplots. This data is also shown in **Fig. 1B.** Error bars, s.d. ($n = 3$)

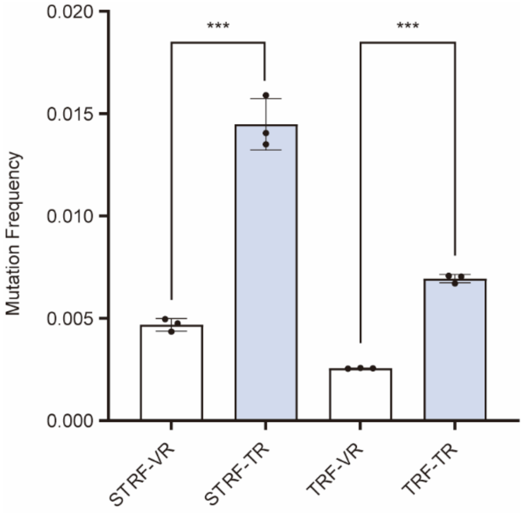


**Figure S7. Mutation frequency (5h) of natural BPP-1 TR shows higher mutation frequency than VR for both short TR (STRF) and long TR (TRF).** The mutation frequencies were calculated from the average of all A position mutations in TR and VR respectively. The natural BBP1 TR and VR sequences were used. Statistical difference was determined by a two-tailed unpaired $t$ test for STRF, $p = 0.0002$, $t = 13.13$, and a two-tailed Welch's $t$ test for TRF, $p = 0.0007$, $t = 36.91$. $p$ value summary: ****$p$ value < 0.0001, 0.0001 < ***$p$ value < 0.001, 0.001 < **$p$ value < 0.01, 0.01 < *$p$ value < 0.05, $p$ value ≥ 0.05: n.s. Error bars, s.d. ($n = 3$)

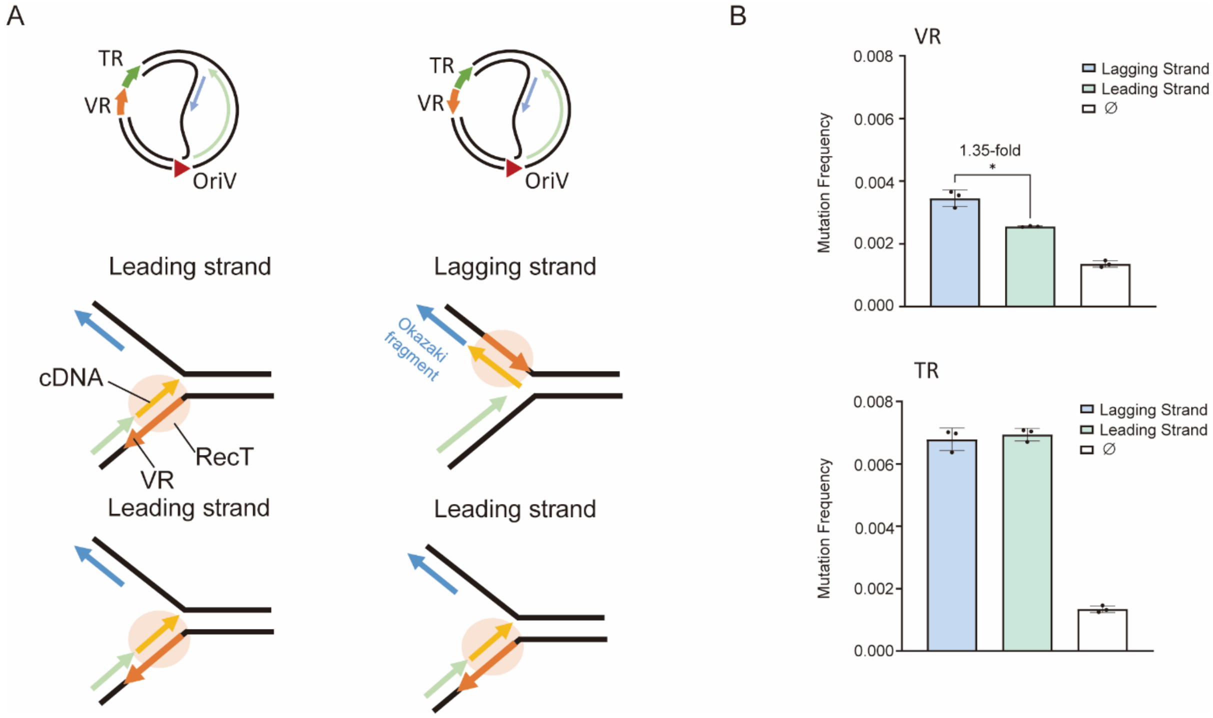


**Figure S8. Optimization of VR orientation relative to the replication fork can improve the mutation frequency. A,** VR orientation relative to the origin of replication determines the leading strand and lagging strand. **B,** Mutation frequency (5h) in VR (top) and TR (bottom) based on the lagging strand vs leading strand targeting. The natural BBP1 TR and VR sequences were used in these experiments. The TR orientation and sequence remained unchanged in these experiments. The mutation frequencies were calculated based on the average of all the A position mutations in TR and VR respectively. Statistical difference was determined by a two-tailed Welch's $t$ test for VR, $p = 0.0276$, $t = 5.839$. $p$ value summary: ****$p$ value < 0.0001, 0.0001 < ***$p$ value < 0.001, 0.001 < **$p$ value < 0.01, 0.01 < *$p$ value < 0.05, $p$ value ≥ 0.05: n.s. Error bars, s.d. ($n = 3$)

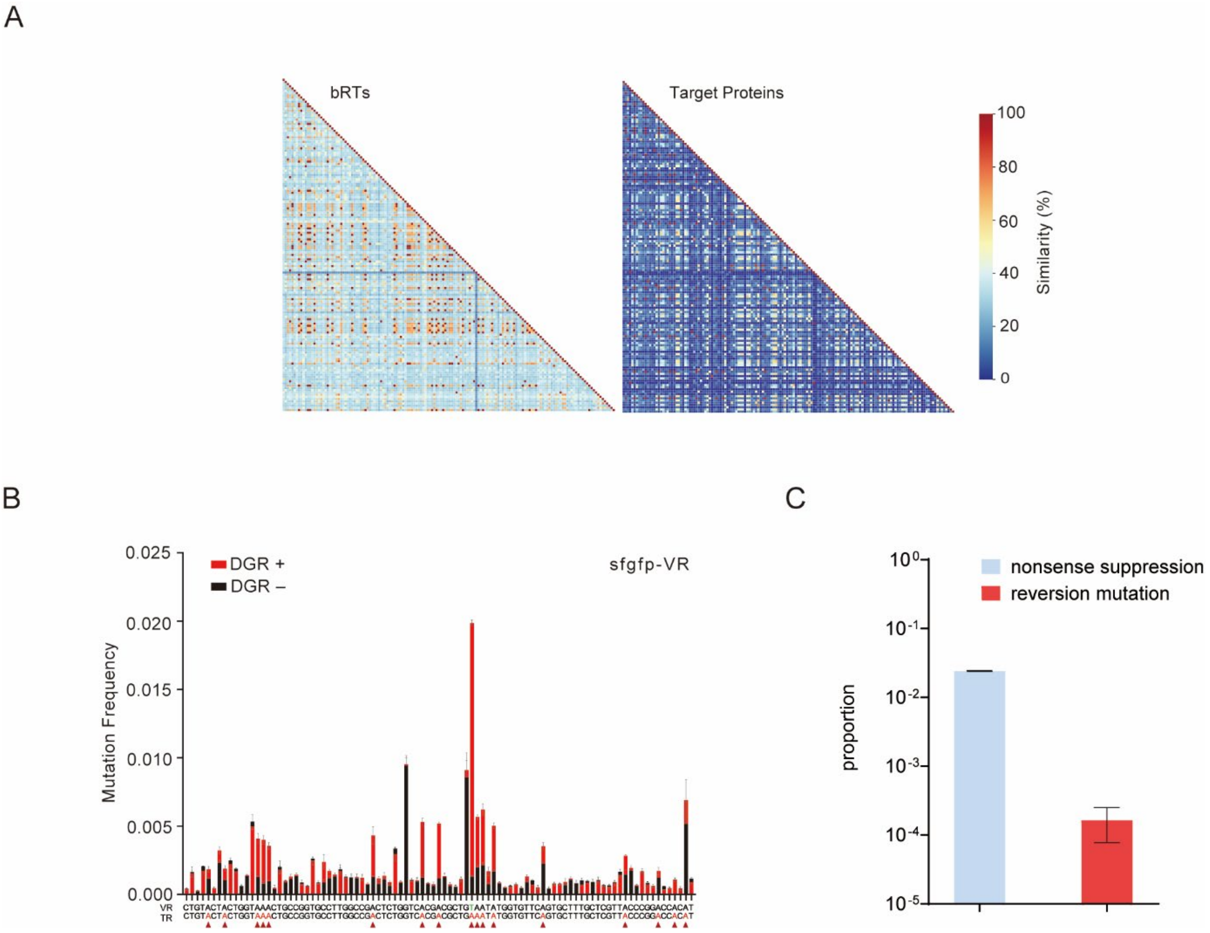


**Figure S9. Expanding VR to a non-WT gene. A,** Similarity matrix computed from multi-sequence alignment of BPP1-like reverse transcriptases (bRTs) and their corresponding target proteins showed that similar bRTs can have diverse target proteins, suggesting VR can be expanded to non-WT proteins. **B,** Mutation frequency barplot of a non-WT gene (*sfgfp*) targeted with a reprogrammed TR. **C,** The proportion of nonsense repression and reversion mutations based on NGS reads. Error bar, s.d. ($n = 3$)

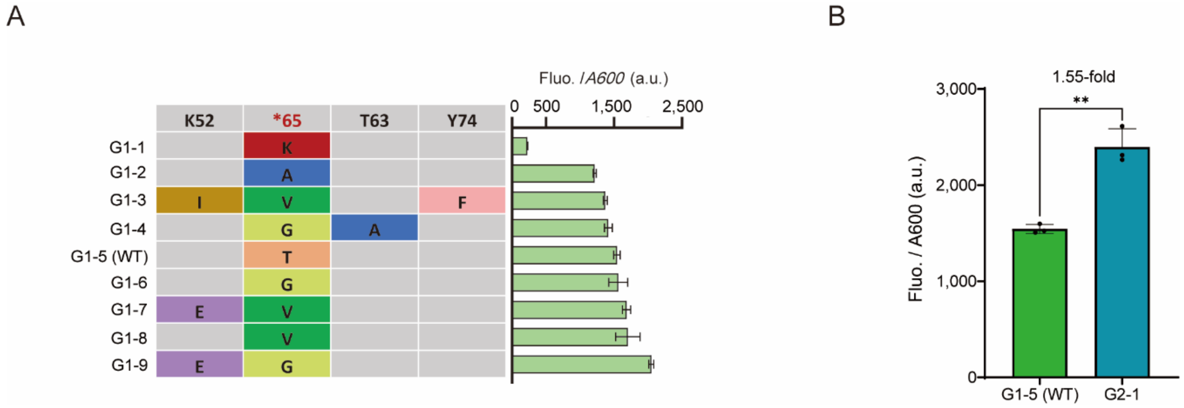


**Figure S10. Fluorescence-activated cell sorting (FACS) enriched multiple sfGFP variants. A,** Results of quantitative fluorescence analysis of single clones selected from a cell population enriched by fluorescence-activated cell sorting (FACS). **B,** In randomly selected colonies, the most fluorescent variant was compared with the original superfolder-GFP. Statistical difference was determined by a two-tailed unpaired *t* test: $p = 0.0016$, $t = 7.558$. *p* value summary: ****$p$ value $< 0.0001$, $0.0001 <$ ***$p$ value $< 0.001$, $0.001 <$ **$p$ value $< 0.01$, $0.01 <$ *$p$ value $< 0.05$, $p$ value $\geq 0.05$: n.s. Error bar, s.d. ($n = 3$)

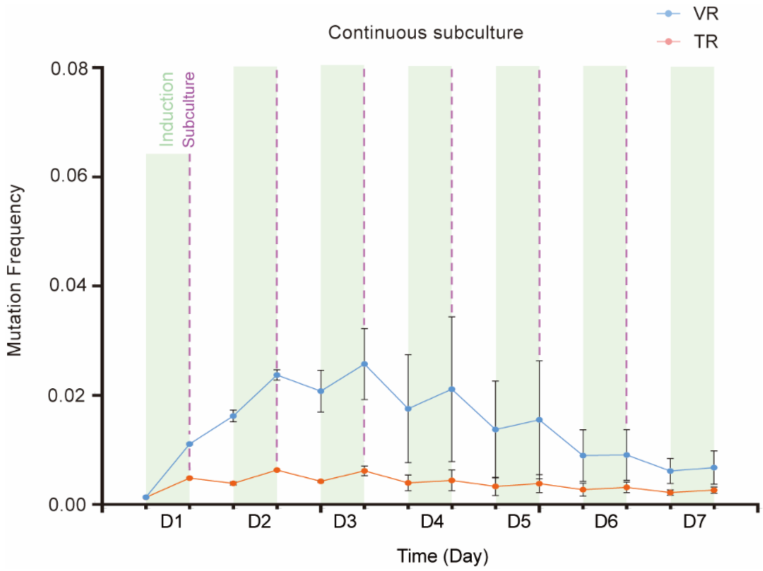


**Figure S11. Mutation frequency of VR and TR of continuous serial subculture of 7 days.** All mutation frequencies (10h) were calculated from the average of all mutations at adenine positions in the corresponding VR or TR. The green blocks represent the induction phases, and the dotted lines represents the subculture steps. The thymine site T193 in VR, which do not match the TR sequence is not included in the calculation. Error bar, s.d. ($n = 3$)

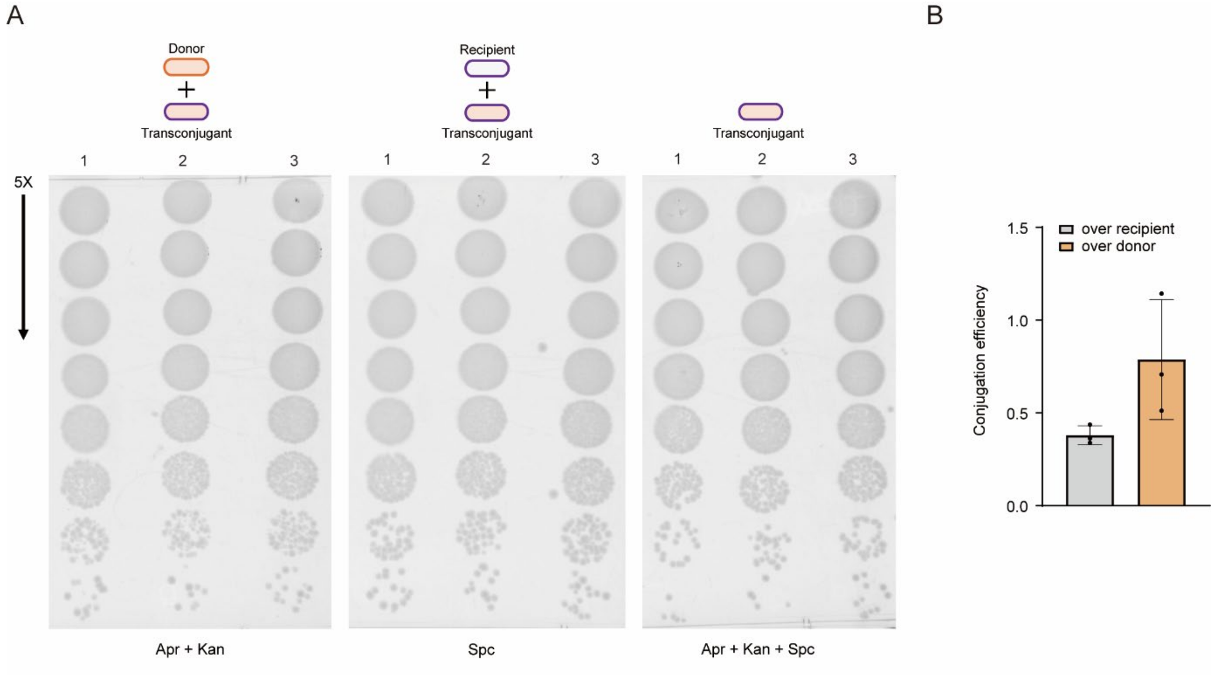


**Figure S12. Quantification of conjugation efficiency of engineered conjugative-plasmid carrying the DGR cassette. A,** Spotting assay for the conjugative-plasmid of DGR conjugation into empty cells as the recipient. The cells were spotted on Apr + Kan (donor), Spec (recipient) and Apr + Kan + Spec (transconjugant) containing agar plates, respectively, with 5-fold serial dilutions. The data for replicate 1 is also shown in **Fig. 3B.** Error bars, s.d. ($n = 3$)

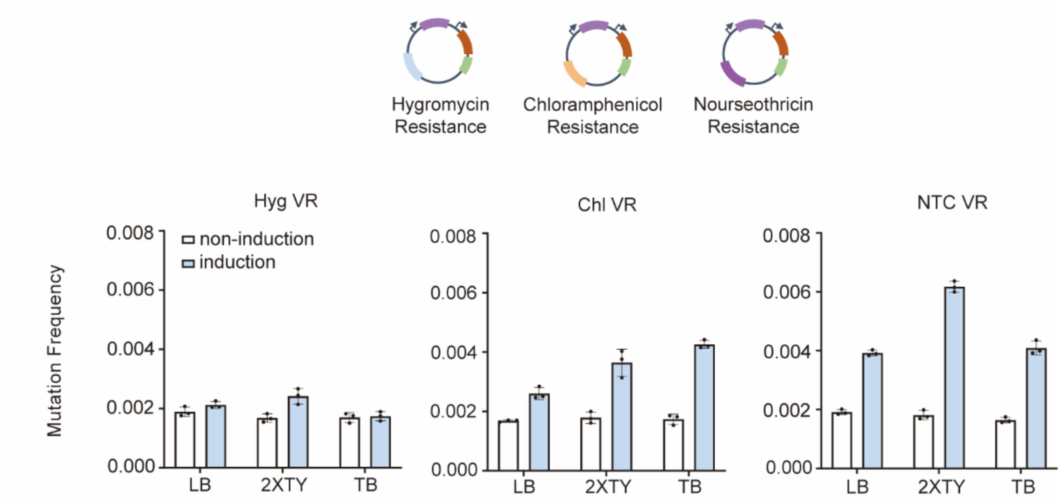


**Figure S13. Mutation Frequency of conjugative-plasmid sfGFP VR for DGR induction in different media (LB, 2XTY, TB) and different resistance genes on the TR plasmid (Hygromycin, Chloramphenicol, Nourseothricin).** All mutation frequencies (10h) were calculated from the average of all A positions mutations in the corresponding VR sequences. Error bars, s.d. ($n = 3$)

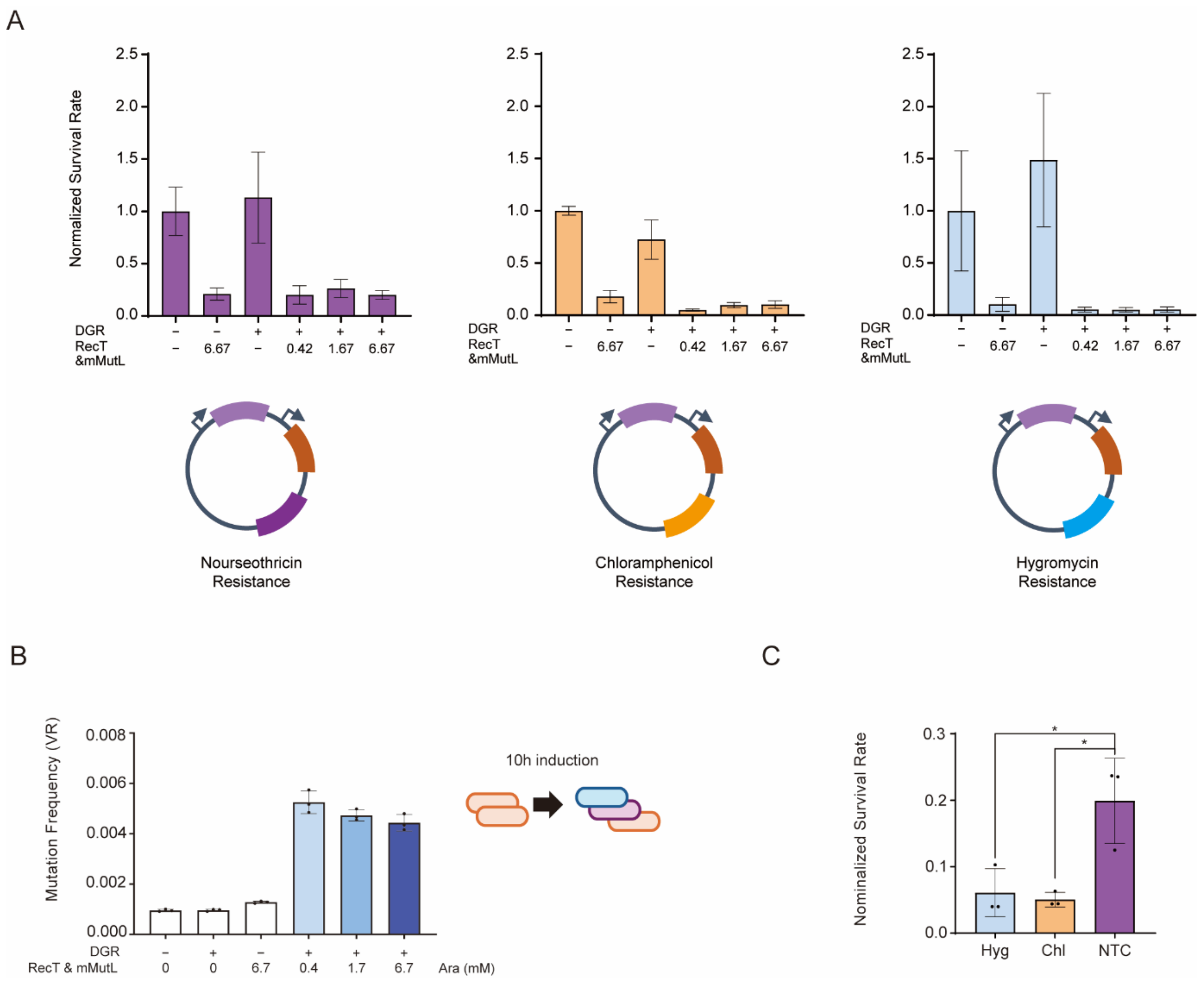


**Figure S14. Optimization of HGT-DGR survival rate. A,** Test of survival rate with different induction levels of CspRecT/mMutL (arabinose, mM) in the context of TR plasmids harbouring different antibiotic resistance genes (hygromycin, chloramphenicol or nourseothricin resistance). **B,** Test of mutation frequency (10h) for each induction level of CspRecT/mMutL. The TR plasmid used harboured nourseothricin resistance. **C,** Survival rate based on TR plasmids harbouring different antibiotic resistance genes (hygromycin, chloramphenicol or nourseothricin resistance) under 0.42 mM arabinose. This data is also shown in **Fig. S12A**. All mutation frequencies were calculated from the average of all A positions mutations in the corresponding VR or TR. Statistical difference was determined by a two-tailed unpaired $t$ test: (Hyg/NTC), $p = 0.0315$, $t = 3.246$, (Chl/NTC), $p = 0.0167$, $t = 3.955$. $p$ value summary: ****$p$ value $< 0.0001$, $0.0001 <$ ***$p$ value $< 0.001$, $0.001 <$ **$p$ value $< 0.01$, $0.01 <$ *$p$ value $< 0.05$, $p$ value $\geq 0.05$: n.s. Error bars, s.d. ($n = 3$)

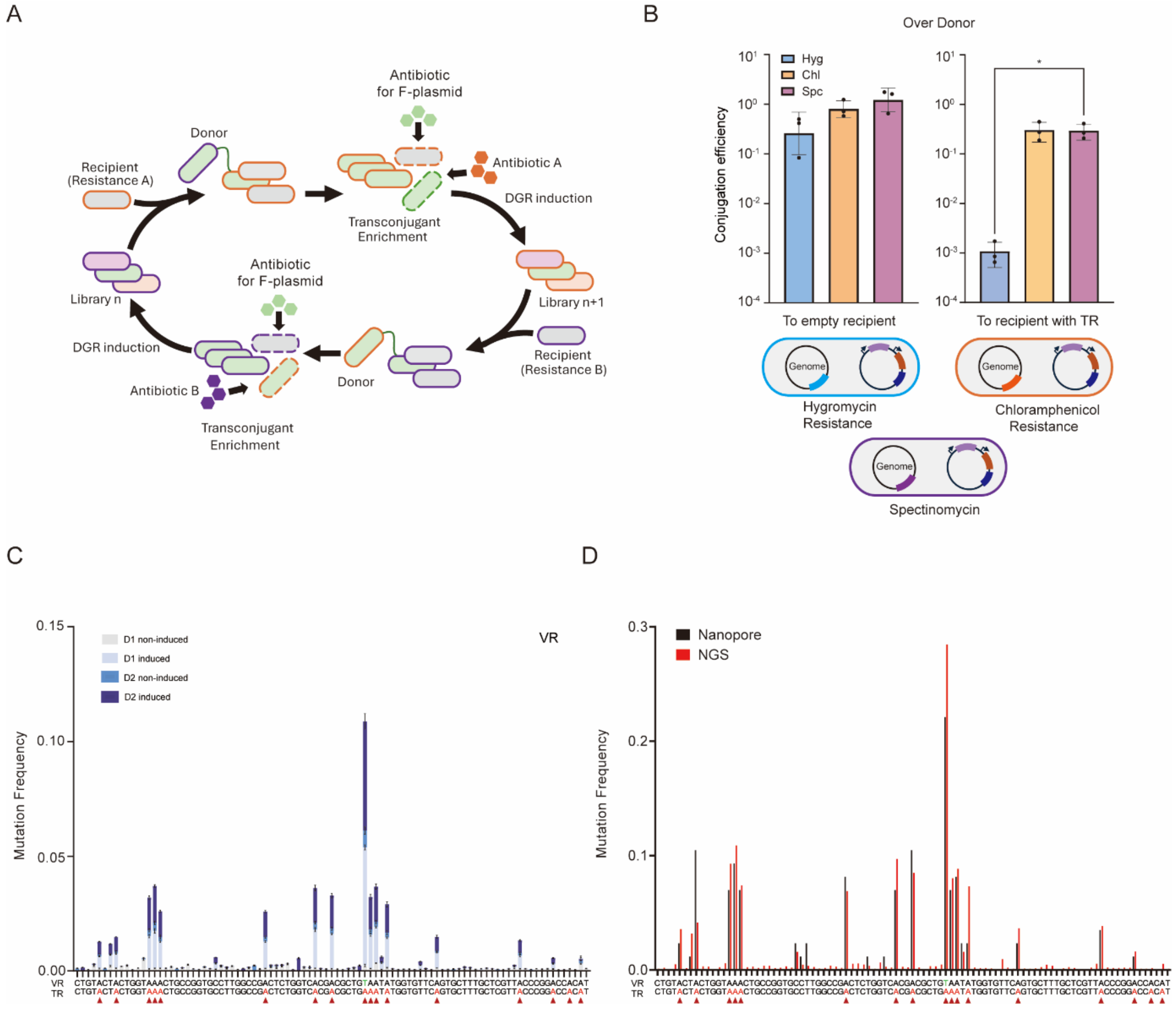


**Figure S15.HGT-DGR enables the transfer of library. A,** Schematic of the full HGT-DGR pipeline where the alternating antibiotics enables the continuous transfer of the library. **B,** Quantification of conjugation efficiency to empty cells or recipient harboring the TR plasmid (Nourseothricin). The recipients contain genomically integrated antibiotic markers (Hygromycin, Chloramphenicol, Spectinomycin). Statistical difference was determined by a two-tailed Welch's $t$ test: (Hyg/Spc), $p = 0.0477$, $t = 4.413$. **C,** Mutational frequency (10h) barplot showed accumulation of mutations over 2-day HGT-DGR (D1: day1, D2: day2). The data of D1 is also shown in **Fig. 3C. D,** A comparison of the mutation profiles obtained from single-colony sequencing and NGS. The samples were subjected to 7 days of conjugation and DGR induction in DH10B and strains with the three exonuclease genes (sbcB, recJ and exoX) knocked out. $p$ value summary: ****$p$ value < 0.0001, 0.0001 < ***$p$ value < 0.001, 0.001 < **$p$ value < 0.01, 0.01 < *$p$ value < 0.05, $p$ value ≥ 0.05: n.s. Error bars, s.d. ($n = 3$)

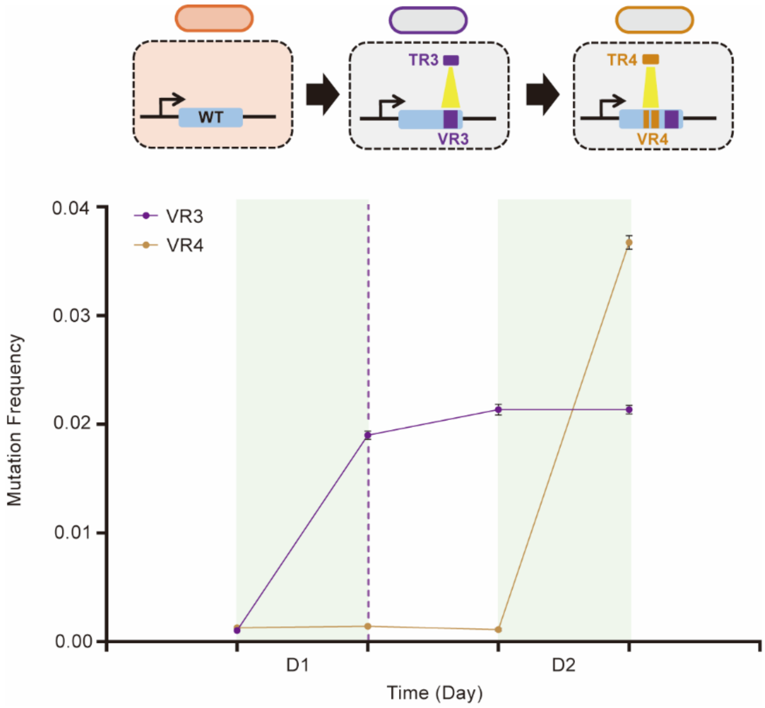


**Figure S16. Rapid replacement of TR and reprogramming of DGR targeting via conjugation.** Mutation frequency (10h) of VR3 (purple line) and VR4 (golden line). All mutation frequencies were calculated from the average of all adenine positions. The green blocks represent the induction phases, and the dotted lines represents the conjugation steps. Error bars, s.d. ($n = 3$)

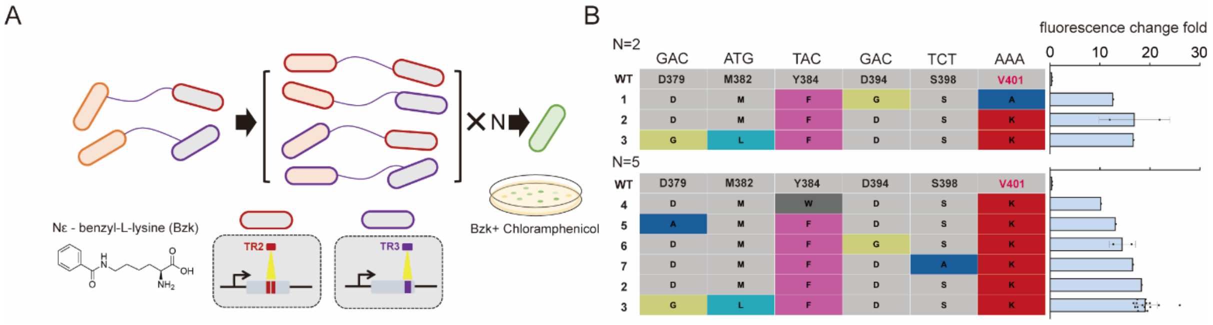


**Figure S17. Evolution of Mm PylRS for incorporation of N$^{\varepsilon}$-Benzylysine (BenzK) via HGT-DGR. A,** Schematic of evolution with pools of TR2 and TR3. **B,** Table of mutations enriched from the BenzK evolution experiment with the dynamic range of the BenzK-dependent expression of GFP shown on the right. The top table shows solutions after 2 rounds of evolution (N = 2) and the bottom table shows solutions after 5 rounds of evolution (N = 5). Error bars, s.d. ($n = 3$)

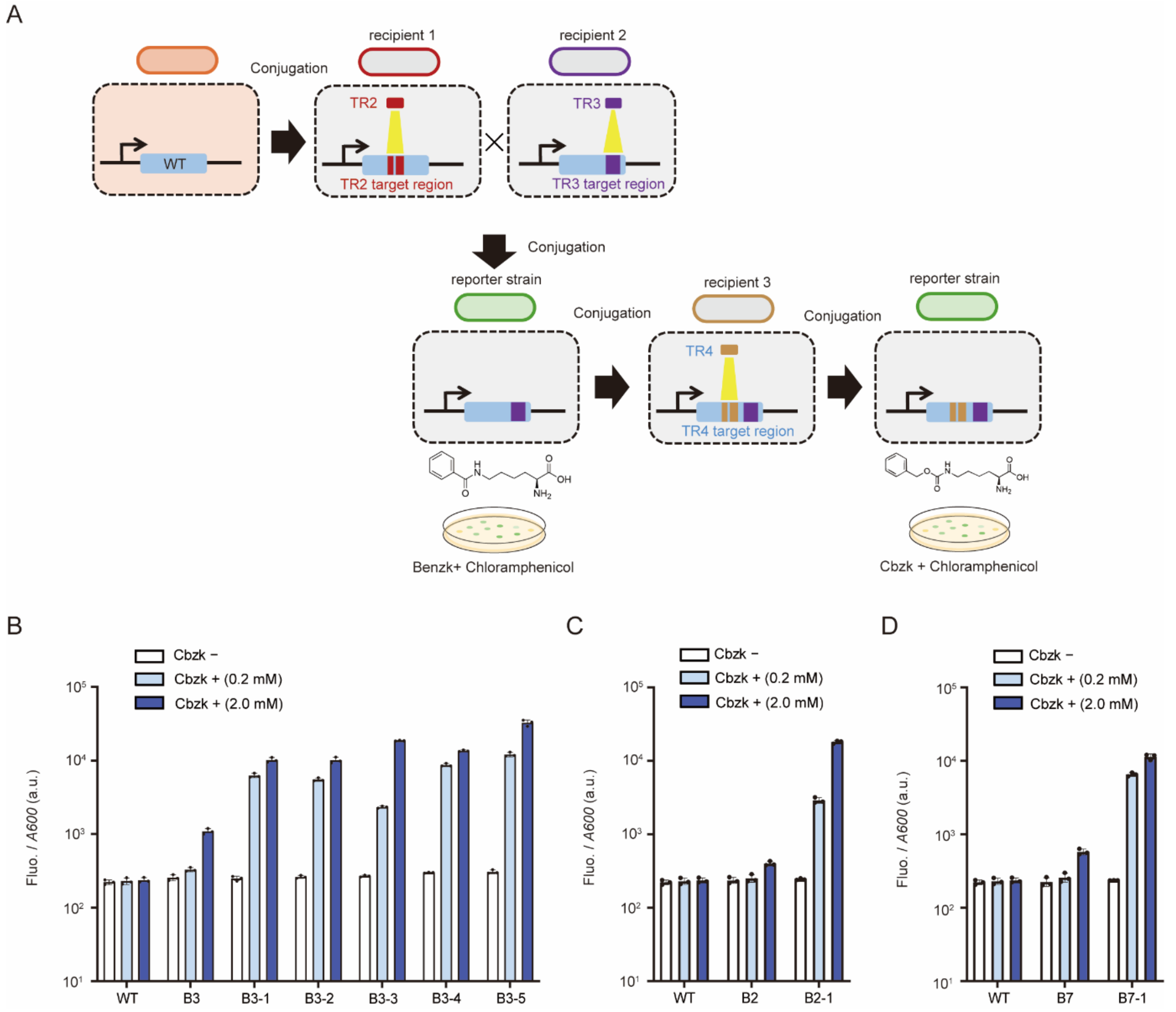


**Figure S18. Evolution of Mm PylRS to expand the substrate scope from Benzk to CbzK. A,** Schematic of the evolution route, where the WT Mm PylRS was first evolved with mixed template of TR2 and TR3, and then selected with BenzK. The enriched clones were then evolved with TR4 via HGT-DGR. **B,** Characterization of CbzK-dependent GFP expression of lineage 3 MmPylRS before evolution (B3) and enriched clones after evolution (B3-1, B3-2, B3-3, B3-4, B3-5). **C,** lineage 2 MmPylRS before evolution (B2) and enriched clones after evolution (B2-1). **D,** lineage B7 MmPylRS before evolution (B7) and enriched clones after evolution (B7-1). WT MmPylRS was used as control. The number before the dash indicates the ancestor clone of this lineage and the number after the dash indicates the progeny clone of this lineage. Error bars, s.d. ($n = 3$)

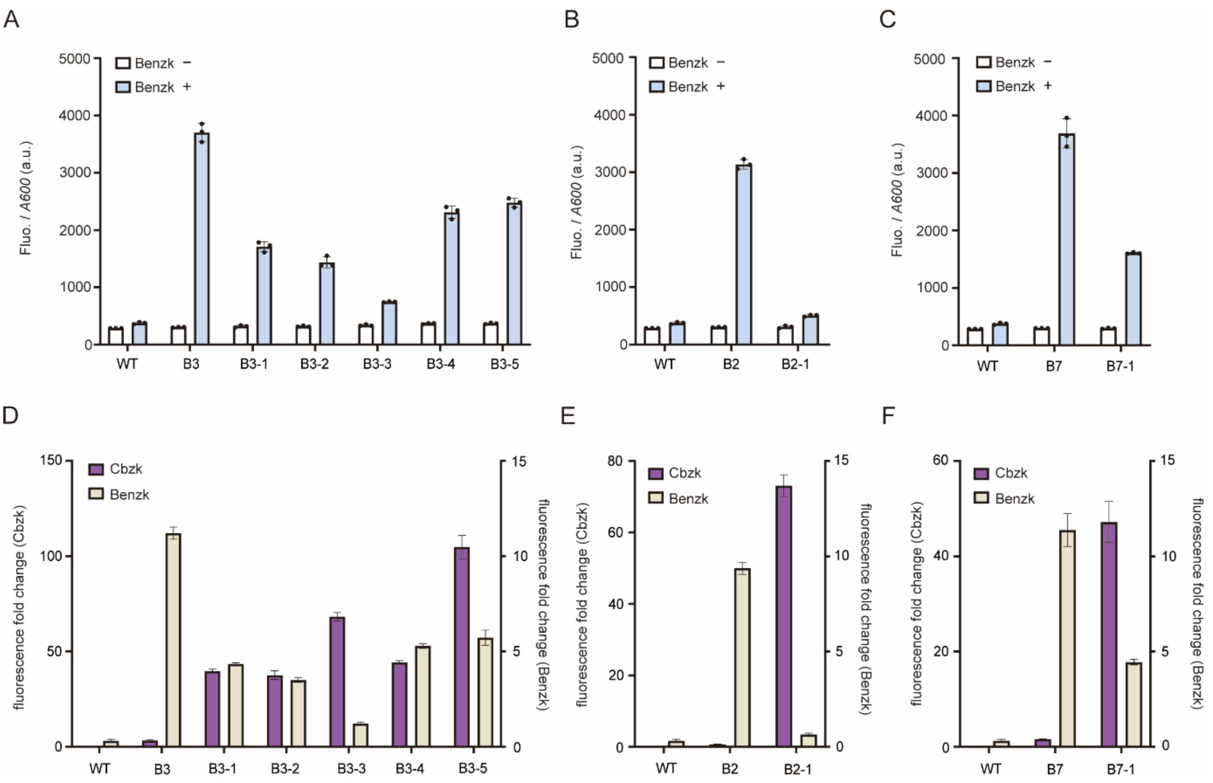


**Figure S19. Characterization of BenzK- and CbzK-dependent GFP expression in the presence of evolved Mm PylRS variants.** **A,** lineage 3 MmPylRS before evolution (3) and enriched clones after evolution (B3-1, B3-2, B3-3, B3-4, B3-5). **B,** lineage B2, MmPylRS before evolution (B2) and enriched clone after evolution (B2-1). **C,** lineage B7, MmPylRS before evolution (B7) and enriched clone after evolution (B7-1). Fluorescence fold change of CbzK-dependent and BenzK-dependent GFP expression. **D,** lineage 3. **E,** lineage 2. **F,** lineage 7. The left y-axis indicates the CbzK fold change and the right Y-axis indicates the BenzK fold change. The data for B3 and B3-5, B2 and B2-1, and B7 and B7-1 are also shown in **Fig. 4F**. Error bars, s.d. ($n = 3$)

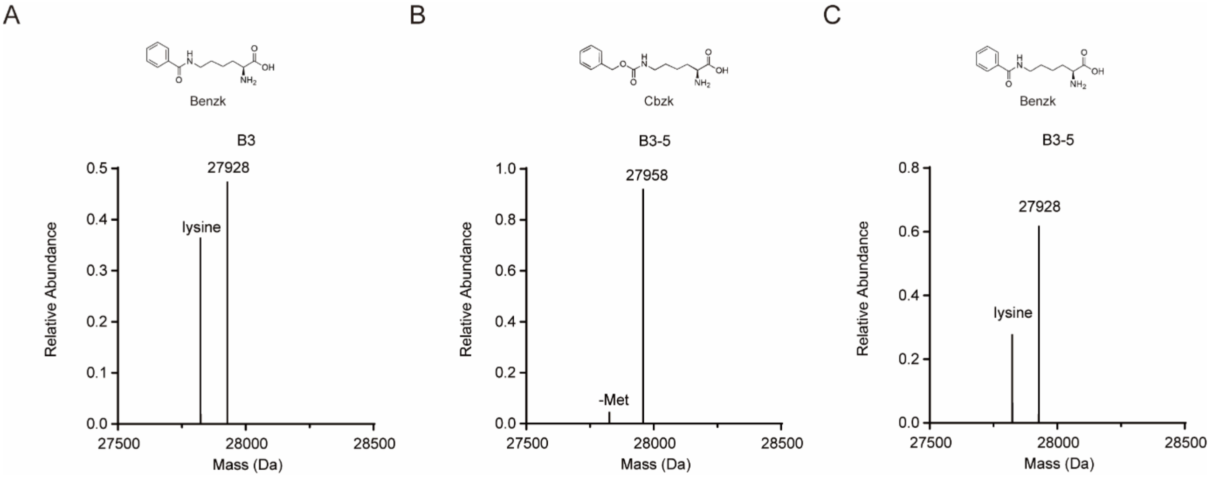


**Figure S20. Mass spectrometry analysis confirmed the ability of variants B3 and B3-5 to utilize Cbzk or Benzk.** A, Mass spectrometry analysis results of variant B3, cultured with Benzk. B, Mass spectrometry analysis results of variant B3-5, cultured with Cbzk. C, Mass spectrometry analysis results of variant B3-5, cultured with Benzk.

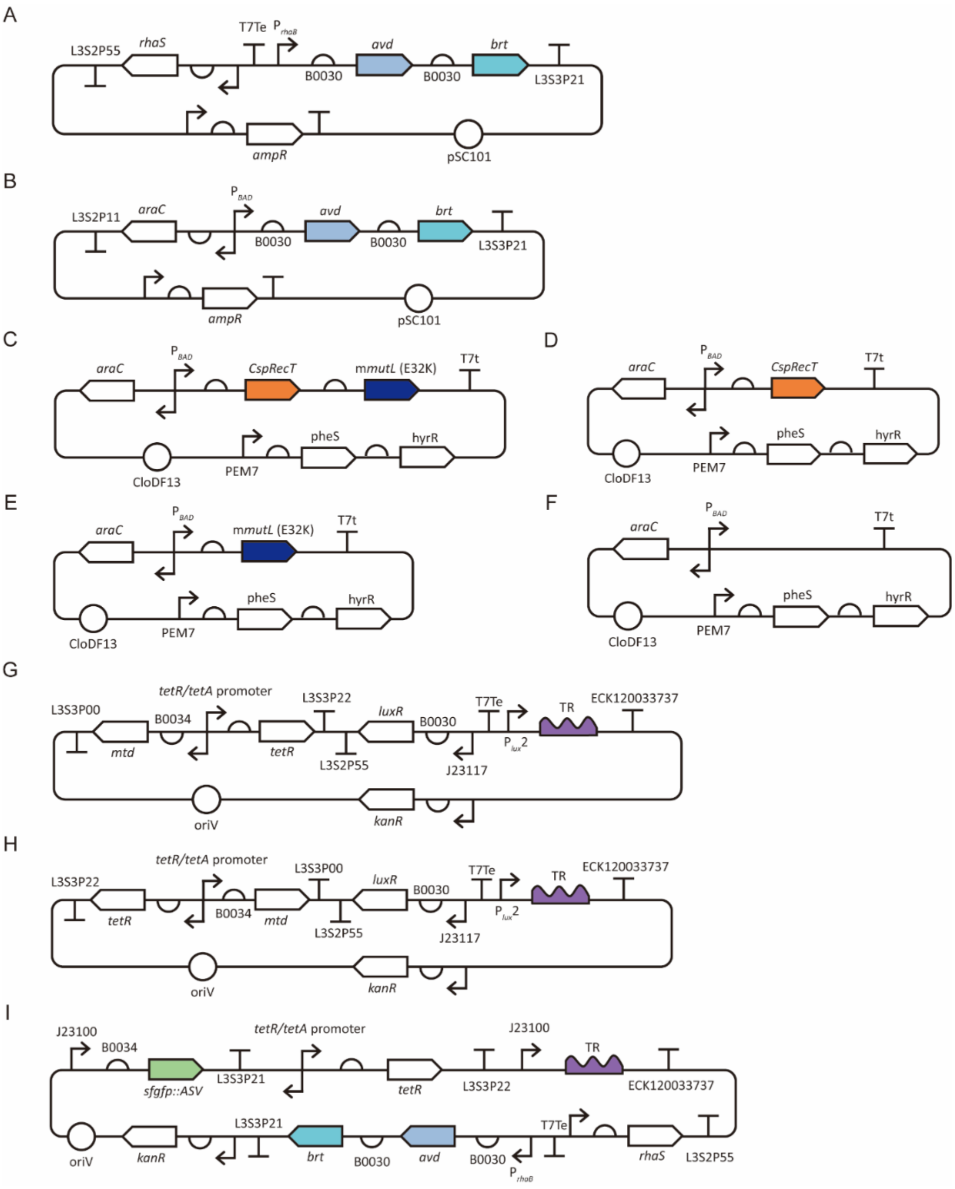


**Figure S21. Representative plasmid maps for key circuits used in the tests of DGR function. A.** The plasmid map of pLG-1 was used in the experiments in **Fig. 1B, C**, and in the tests of **Fig. S5** and **Fig. S6**. **B.** The plasmid map of pLG-2 was used in the experiments in **Fig. S4**, **Fig. S7** and **Fig. S8**. **C.** The plasmid map of pFR157 was used in the experiments in **Fig. 1B, C**, **Fig. 2A, B**, and in the tests of **Fig. S4**, **Fig. S5**, **Fig. S6**, **Fig. S7**, **Fig. S8**, and **Fig. S9B**. **D.** The plasmid map of pLG-3 was used in the experiments in **Fig. S5** and **Fig. S6**. **E.** The plasmid map of pLG-4 was used in the experiments in **Fig. S5** and **Fig. S6**. **F.** The plasmid map of pLG-5 was used in the experiments in **Fig. S5** and **Fig. S6**. **G.** The plasmid map of pLG-6 and pLG-7. The pLG-6 was used in the experiments in **Fig. 1B. C**, **Fig. S5** and **Fig. S6**, the pLG-7 was used in the experiments in **Fig. S8**. **H.** The plasmid map of pLG-8 and pLG-9, pLG-8 was used in the experiments in **Fig. S4** and **Fig. S7**. pLG-9 was used in the experiments in **Fig. S4**, **Fig. S7**, and **Fig. S6**. **I.** The plasmid map of pLG-10, pLG-11 and pLG-12, pLG-10 was used in the experiments in **Fig. 2A**, **Fig. 2B**, and **Fig. S9B**. pLG-11 and pLG-12 were used in the tests in **Fig. 2A**.

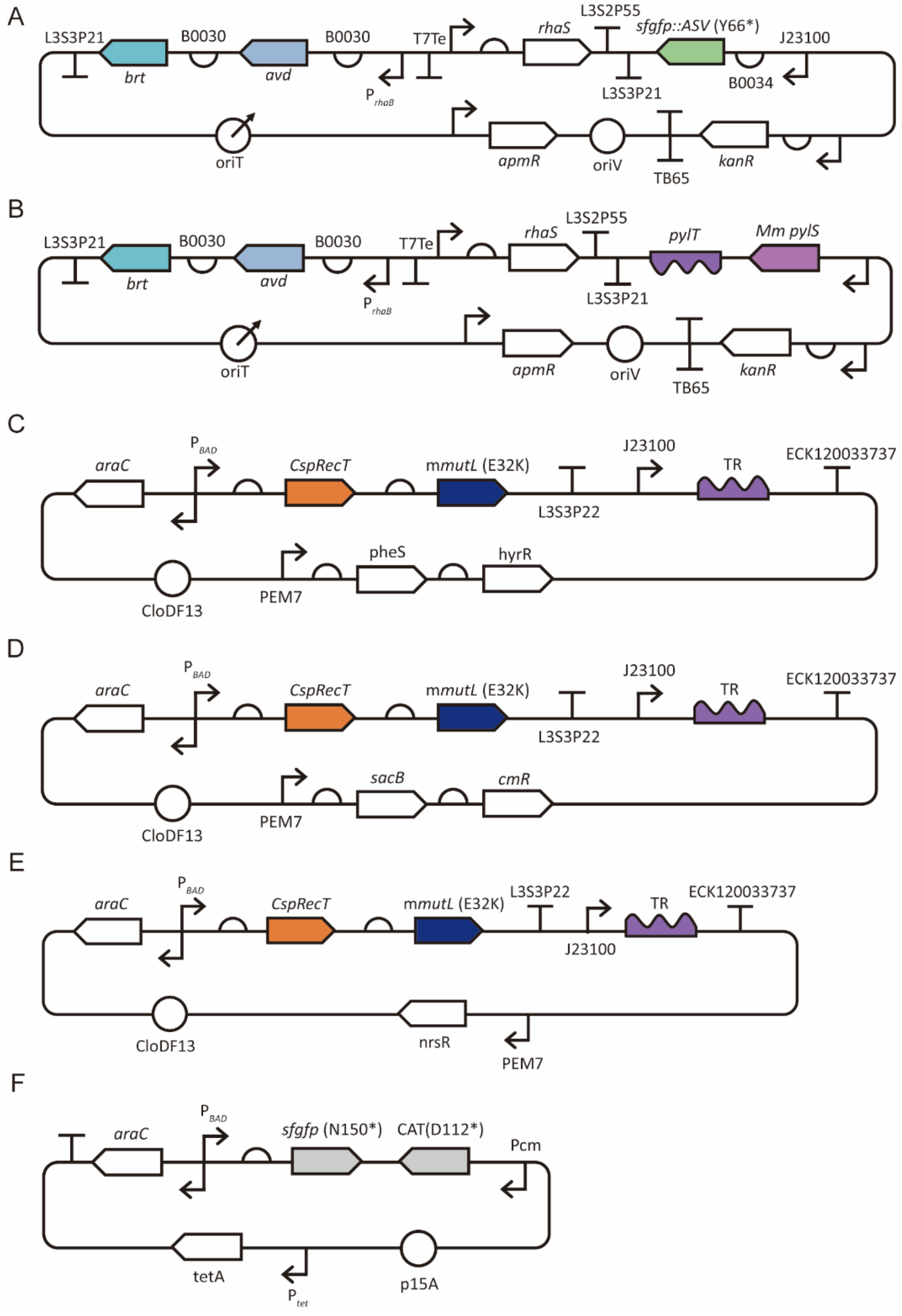


**Figure S22. Representative plasmid maps for key constructs used in the HGT-mediated DGR. A.** The plasmid map of pLG-13 was used in the experiments in **Fig. 3C, D**, and in the tests of **Fig. S11** – **Fig. S15**. **B.** The plasmid map of pLG-14 was used in the experiments in **Fig. 4**, and **Fig. S16** – **Fig. S20**. **C.** The plasmid map of pLG-15 was used in the experiments in **Fig. S13, Fig. S14A**, and **Fig. S14C**. **D.** The plasmid map of pLG-16 was used in the experiments in **Fig. S13, Fig. S14A**, and **Fig. S14C**. **E.** The plasmid map of pLG-17 – pLG-21. pLG-17 was used in the experiments in **Fig. 3**. and was used in the tests in **Fig. S13** – **Fig. S15**. pLG-18 was used in the experiments in **Fig. 4B, D**. pLG-19 and pLG-20 were used in the experiments in **Fig. 4E, F**, and were used in the tests in **Fig. S16** – **Fig. S20**. pLG-21 was used in the experiments in **Fig. 4E, F**, and was used in the tests in **Fig. S17** and **Fig. S18**. **F.** The plasmid map of pFR361, was used in the experiments in **Fig. 4**, and was used in the tests **in Fig. S17** – **Fig. S20**.

**Table S1. Plasmids involved in this study**

| No. | Name | Function | Resource |
|---|---|---|---|
| 1 | pLG-1 | Rhamnose induced DGR device (*avd* & *brt*) | This study |
| 2 | pLG-2 | Arabinose induced DGR device (*avd* & *brt*) | This study |
| 3 | pFR157 | Arabinose induced m*mutL* and *CsprecT* generator | This study |
| 4 | pLG-3 | Arabinose induced *CsprecT* generator | This study |
| 5 | pLG-4 | Arabinose induced m*mutL* generator | This study |
| 6 | pLG-5 | Negative control for pFR157, pLG-3, and pLG4 | This study |
| 7 | pLG-6 | reversed WT VR and AHL induced WT TR (short) | This study |
| 8 | pLG-7 | reversed WT VR and AHL induced WT TR | This study |
| 9 | pLG-8 | WT VR and AHL induced WT TR (short) | This study |
| 10 | pLG-9 | WT VR and AHL induced WT TR | This study |
| 11 | pLG-10 | Rhamnose induced DGR device (*avd* & *brt*), with WT VR and consititutively expressed WT TR (93 bp) | This study |
| 12 | pLG-11 | Rhamnose induced DGR device (*avd* & *brt*), with WT VR and consititutively expressed WT TR (60 bp) | This study |
| 13 | pLG-12 | Rhamnose induced DGR device (*avd* & *brt*), with WT VR and consititutively expressed WT TR (80 bp) | This study |
| 14 | pLG-13 | Birmingham IncP alpha plasmid carrying Rhamnose induced DGR device and muttated *sfgfp* as VR | This study |
| 15 | pLG-14 | Birmingham IncP alpha plasmid carrying Rhamnose induced DGR device and Mm PylRS | This study |
| 16 | pLG-15 | Arabinose induced m*mutL* and *CsprecT* generator with TR for *sfgfp*, hygromycin resistance | This study |
| 17 | pLG-16 | Arabinose induced m*mutL* and *CsprecT* generator with TR for *sfgfp*, chloramphenicol resistance | This study |
| 18 | pLG-17 | Arabinose induced m*mutL* and *CsprecT* generator with TR for *sfgfp*, nourseothricin resistance | This study |
| 19 | pLG-18 | Arabinose induced m*mutL* and *CsprecT* generator with TR1 for *Mm PylR,* nourseothricin resistance | This study |
| 20 | pLG-19 | Arabinose induced m*mutL* and *CsprecT* generator with TR2 for *Mm PylR,* nourseothricin resistance | This study |
| 21 | pLG-20 | Arabinose induced m*mutL* and *CsprecT* generator with TR3 for *Mm PylR,* nourseothricin resistance | This study |
| 22 | pLG-21 | Arabinose induced m*mutL* and *CsprecT* generator with TR4 for *Mm PylR,* nourseothricin resistance | This study |
| 23 | p15A-$CAT^{112TAG}$-$GFP^{150TAG}$) | Reporter circuit for the Mm PylR evolution | Daniele Cervettini et al. (*48*) |

**Table S2. Strains used in this study**

| ***E. coli* strain** | **Description and genotype** | **Resource** |
|---|---|---|
| DH10b | *Δ(ara-leu) 7697 araD139 fhuA ΔlacX74 galK16 galE15 e14- ϕ80dlacZΔM15 recA1 relA1 endA1 nupG rpsL (StrR) rph spoT1 Δ(mrr-hsdRMS-mcrBC)* derived from *E. coli K-12* | New England BioLabs |
| sLG001 | DH10b Δ*recJ*, Δ*sbcB*, Δ*exoX* for triple knockouts for improved mutagenesis efficiency | This study |
| sLG002 | DH10b Δ*recJ*, Δ*sbcB*, Δ*exoX* + smR, triple KO with genomically integrated spectinomycin resistance for conjugation experiment | This study |
| sLG003 | DH10b Δ*recJ*, Δ*sbcB*, Δ*exoX* + *cmR*, triple KO with genomically integrated chloramphenicol resistance gene for conjugation experiment | This study |
| sLG004 | DH10b Δ*recJ*, Δ*sbcB*, Δ*exoX* + *hygR*, triple KO with genomically integrated chloramphenicol resistance gene for conjugation experiment | This study |

**Table S3. Primers used in this study**

| No. | Name | Sequence | Function |
|---|---|---|---|
| 1 | LG_1_F | GATCTAAAGAGGAGAAAGGATCTGGCACCATC | Forward primers for pLG-6/pLG-7/pLG-8/pLG-9 VR |
| 2 | LG_1_R | GCAATTTTAACAGGAGCAATTGATTGCCCATAC | Reverse primers for pLG-8/pLG-9 VR |
| 3 | LG_2_R | GAATTGATCAACGTCTCATTTTCGCCAGATATC | Reverse primers for pLG-7 VR |
| 4 | LG_3_R | GAATTGATCAACGTCTCATTTTCGCCAGATATC | Reveser primer for pLG-6 VR |
| 5 | LG_4_F | CCTGTAGGATCGTACAGGTTTACGCAAG | Forward primers for pLG-6/pLG-7/pLG-8/pLG-9 TR |
| 6 | LG_4_R | GATAGCGGCGAAAACATTGTGGATGC | Reverse primers for pLG-6/pLG-7/pLG-8/pLG-9 TR |
| 7 | LG_5_F | CGTAAAGGCGAGGAGCTGTTCAC | Forward primers for pLG-10/ pLG-11/ pLG-12 VR |
| 8 | LG_5_R | CCTCATTAAGCAGCTCTAATGCGCTG | Reverse primers for pLG-10/ pLG-11/ pLG-12 VR |
| 9 | LG_6_F | GCATTATATGCACTCAGCGCTGTGG | Forward primers for pLG-10/ pLG-11/ pLG-12 TR |
| 10 | LG_6_R | GCGACAGTAACCACTTTTCGACGC | Reverse primers for pLG-10/ pLG-11/ pLG-12 TR and pLG-13/pLG14-VR |
| 11 | LG_7_F | CTCTACTACGGTTGCCTTGGCGAAGAGCTGCAG | Forward primer for pLG-13 VR |
| 12 | LG_8_F | AACACCCCTTGTATTACTGTTTATGTAAGC | Forward primer for pLG-14 VR |
| 13 | LG_9_F | TAAATTGGCGATGAGAGAGGAGGATTACAAAATGCC | Forward primer for pLG-15/pLG-16/pLG-17/pLG-18/pLG-19/pLG-20/pLG-21 |
| 14 | LG_9_R | TGTCTAACAATTCGTTCAAGCCGAGGGGCC | Reverse primer for pLG-15/pLG-16/pLG-17/pLG-18/pLG-19/pLG-20/pLG-21 |

**Table S4. VR and TR sequences in this study**

| No. | Name | Sequence | Function |
|---|---|---|---|
| 1 | WT-VR | CGCTGCTGCGCTATTCGGCGGCTCCTGGTTCTACA CGTCGTACTCGGGTTCTCGCGCTGCGTACTGGAAC GCCGGGCCGTCGAACTCCAGCGCGAACAT | |
| 2 | WT-TR | CGCTGCTGCGCTATTCGGCGGCAACTGGAACAAC ACGTCGAACTCGGGTTCTCGCGCTGCGAACTGGA ACAACGGGCCGTCGAACTCGAACGCGAACAT | |
| 3 | GFP-VR61 | CTGCCGGTGCCTTGGCCGACTCTGGTCACGACGCT GTAATATGGTGTTCAGTGCTTTGCTC | |
| 4 | GFP-TR61 | CTGCCGGTGCCTTGGCCGACTCTGGTCACGACGCT GAAATATGGTGTTCAGTGCTTTGCTC | |
| 5 | GFP-VR81 | TACTGGTAAACTGCCGGTGCCTTGGCCGACTCTGG TCACGACGCTGTAATATGGTGTTCAGTGCTTTGCT CGTTACCCGGA | |
| 6 | GFP-TR81 | TACTGGTAAACTGCCGGTGCCTTGGCCGACTCTGG TCACGACGCTGAAATATGGTGTTCAGTGCTTTGCT CGTTACCCGGA | |
| 7 | GFP-VR93 | CTGTACTACTGGTAAACTGCCGGTGCCTTGGCCGA CTCTGGTCACGACGCTGTAATATGGTGTTCAGTGC TTTGCTCGTTACCCGGACCACAT | |
| 8 | GFP-TR93 | CTGTACTACTGGTAAACTGCCGGTGCCTTGGCCGA CTCTGGTCACGACGCTGAAATATGGTGTTCAGTGC TTTGCTCGTTACCCGGACCACAT | |
| 9 | VR1 | CGAAGATTTTGATCGGGTCCGGCAGAGCACGGTC CAGTTTACGCAGGTAGTTGTACAGGTTCGGAGCCA GCATCGGACGCAGGCAGAAG | |
| 10 | TR1 | CTTCTGCCTGCGTCCGATGCTGGCTCCGAACCTGA AAAACTACAAACGTAAACTGGACCGTGCTCTGCC GGACCCGATCAAAATCTTCG | |
| 11 | VR2 | CCAGGTTTTCACGGGTGCAACCAGAACCCATCTGG CAGAAGTTCAGCATGGTGAATTCTTCCAGGTGTTC TTTACCGTCAGATTCTTTACGG | |
| 12 | TR2 | CCGTAAAGAATCTGACGGTAAAGAACACCTGGAA GAATTCACCATGCTGAACTTCAATCAGATGGGTTC TGGTTGCACCCGTGAAAACCTGG | |
| 13 | VR3 | CACGGTCCAGCGGGATCGGACCAACAACAGCAGA AGACAGTTCCAGGTCACCGTGCATAACGTCCAGG GTGTCACCGTAAACCATGCAAGAGTCACCAACGA | |
| 14 | TR3 | TCGTTGGTGACTCTTGCATGGTTTACGGTGACACC CTGGACGTTATGCACGGTGACCTGGAACTGTCTTC TGCTAAAGTTGGTCCGATCCCGCTGGACCGTG | |
| 15 | VR4 | CACGGGTGCAACCAGAACCCATCTGGCAGAAGTT CAGCATGGTGAATTCTTCCAGGTGTTCTTTACCGTC AGATTCTTTACGGTAGCACGGACCG | |
| 16 | TR4 | CGGTCCGTGCTACCGTAAAGAATCTGACGGTAAAG AACACCTGGAAGAATTCACCAAGCTGAACTTCAA CCAGATGGGTTCTGGTTGCACCCGTG | |
